# Integrated transcriptomic landscape of medulloblastoma and ependymoma reveals novel tumor subtype-specific biology

**DOI:** 10.1101/2024.10.21.619495

**Authors:** Sonali Arora, Nicholas Nuechterlein, Matt Jensen, Gregory Glatzer, Philipp Sievers, Srinidhi Varadharajan, Andrey Korshunov, Felix Sahm, Stephen C. Mack, Michael D. Taylor, Taran Gujral, Eric C Holland

## Abstract

Medulloblastoma and ependymoma are common pediatric central nervous system tumors with significant molecular and clinical heterogeneity. We collected bulk RNA sequencing data from 888 medulloblastoma and 370 ependymoma tumors to establish a comprehensive reference landscape. Following rigorous batch effect correction, normalization, and dimensionality reduction, we constructed a unified landscape to explore gene expression, signaling pathways, RNA fusions, and copy number variations. Our analysis revealed distinct clustering patterns, including two primary ependymoma compartments, EPN-E1 and EPN-E2, each with specific RNA fusions and molecular signatures. In medulloblastoma, we observed precise stratification of Group 3/4 tumors by subtype and in SHH tumors by patient age. This landscape serves as a vital resource for identifying biomarkers, refining diagnoses, and enables the mapping of new patients’ bulk RNA-seq data onto the reference framework to predict biology and outcome from nearest neighbor analysis facilitate accurate disease subtype identification. The landscape is accessible via Oncoscape, an interactive platform, empowering global exploration and application.

**One Sentence Summary:** A landscape built using only Transcriptomic analysis for medulloblastoma and ependymoma reveals novel insights about subtype-specific biology.

## INTRODUCTION

Medulloblastoma, a highly malignant primary brain tumor originating in the cerebellum, is the most common pediatric central nervous system cancer, accounting for nearly 20% of all childhood brain tumors^1^. Historically considered a single disease entity, medulloblastoma is now understood to encompass four distinct molecular subtypes as identified in the WHO 2021 classification: Wingless-type (Wnt), Sonic hedgehog (SHH), Group 3, and Group 4.^2^ These subtypes differ not only in their molecular characteristics but also in their clinical behavior, prognosis, and response to treatment. Advances in genomic technologies have revealed a complex landscape of genetic mutations, copy number variations, and epigenetic alterations across these subtypes, deepening our understanding of tumor biology and uncovering potential targets for precision medicine.

Ependymomas (EPNs) are tumors of the neuroepithelial cells, presenting across all age groups and occurring at various locations along the central nervous system. In pediatric populations, ependymomas represent about 10% of all malignant central nervous system tumors, with a significant portion (30%) diagnosed in children younger than three years^3^. Recent advancements in DNA methylation and gene expression profiling have allowed for the identification of distinct molecular subtypes of ependymomas, each with unique clinical and pathological features. In the supratentorial region, ependymomas are characterized by two primary molecular subtypes driven by recurrent gene fusions: one involving the ZFTA gene (previously known as C11orf95, often fused with RELA), and another involving YAP1^4,5^. In the posterior fossa, ependymomas are now classified into two molecular subtypes, PF-A and PF-B, with an additional classification for PF NEC/NOS tumors. These molecular distinctions have important implications for the diagnosis, treatment, and prognosis of patients with ependymoma, underscoring the necessity for precise molecular characterization in clinical practice.

In this study, we present a comprehensive visual integration method for analyzing bulk RNA seq from a large cohort of medulloblastoma and ependymoma cases. We harmonized and integrated transcriptional data from five publicly accessible medulloblastoma studies^6–9^ and eight publicly accessible ependymoma studies. After correcting for batch effects and normalizing the data, we employed dimensionality reduction techniques to construct a reference landscape that reveals significant patterns within this aggregated multi-disease dataset. While the previously published analysis^10^ which aids in visualizing and analyzing different brain diseases, contained ependymoma and medulloblastoma patient samples, the number of samples were limited to only those derived from Children Brain Tumor Network (93 and 117 samples respectively). This represents the largest collection of bulk RNA-seq profiles for medulloblastoma and ependymoma.

Transcriptomic landscape analysis provides several critical insights and capabilities. Our analysis recapitulates known molecular subtypes in medulloblastoma but also enables detailed examination of alterations in gene expression, signaling pathways, gene fusions, and copy number profiles across these tumors.

This landscape analysis facilitates the visualization of both common and unique features across different tumor subtypes, additionally it also aids in identifying potential misdiagnoses and guiding treatment decisions for new patients. Furthermore, our inclusion of fetal brain samples allows for a deeper understanding of the similarities of these tumors to specific CNS developmental stages.

Finally, by making this resource available through the interactive platform Oncoscape^11^ (https://oncoscape.sttrcancer.org/medepn2024.html) we empower researchers and clinicians to explore genes of interest, discover novel biomarkers, and accelerate the pace of research and discovery in the field of neuro-oncology.

## RESULTS

### Constructing a reference landscape for medulloblastoma and ependymoma

We gathered 370 ependymoma samples and 888 medulloblastoma samples from North America and Europe to construct a comprehensive reference landscape for both tumor types. The ependymoma cohort^4,12–15^ included 134 supratentorial, 135 posterior fossa, 77 ependymoma (NOS), 11 anaplastic, 9 myxopapillary, and 4 spinal ependymoma samples, sourced from across North America and Europe. The medulloblastoma cohort^7,16^ consisted of 364 Group 4, 229 Group 3, 274 SHH, 9 WNT, and 12 medulloblastoma (NOS) samples, all collected from North America. Additionally, we incorporated 100 healthy brain samples at various stages during embryonic and post-natal development^17^ to serve as a control dataset. These control samples comprised 48 forebrain and 52 hindbrain samples, covering a broad developmental range: 65 samples were from 4 to 19 weeks post-conception and 35 post-natal samples (**Fig S1**).

Raw sequencing reads from each sample were aligned to the human genome reference hg38, and gene counts were obtained for each gene. Focusing on protein-coding genes, we corrected for batch effects using the ComBatSeq function from the R package “sva”. Gene expression data was then normalized using variance stabilizing transformation (VST). To ensure the robustness of the clustering, we performed a comprehensive comparison of Principal component analysis (PCA), t-distributed stochastic neighbor embedding (t-SNE), and uniform manifold approximation and projection (UMAP) with and without batch correction and explored various normalization methods **(Fig S1a-i).**^10,18^.

After overlaying known biological information, we selected the VST-normalized UMAP as our final reference landscape because it demonstrated no batch effects based on data source **(Fig. 1a)** and effectively captured clusters corresponding to publicly known subtypes of the disease (**Fig. 1b**).

**Fig. 1.**
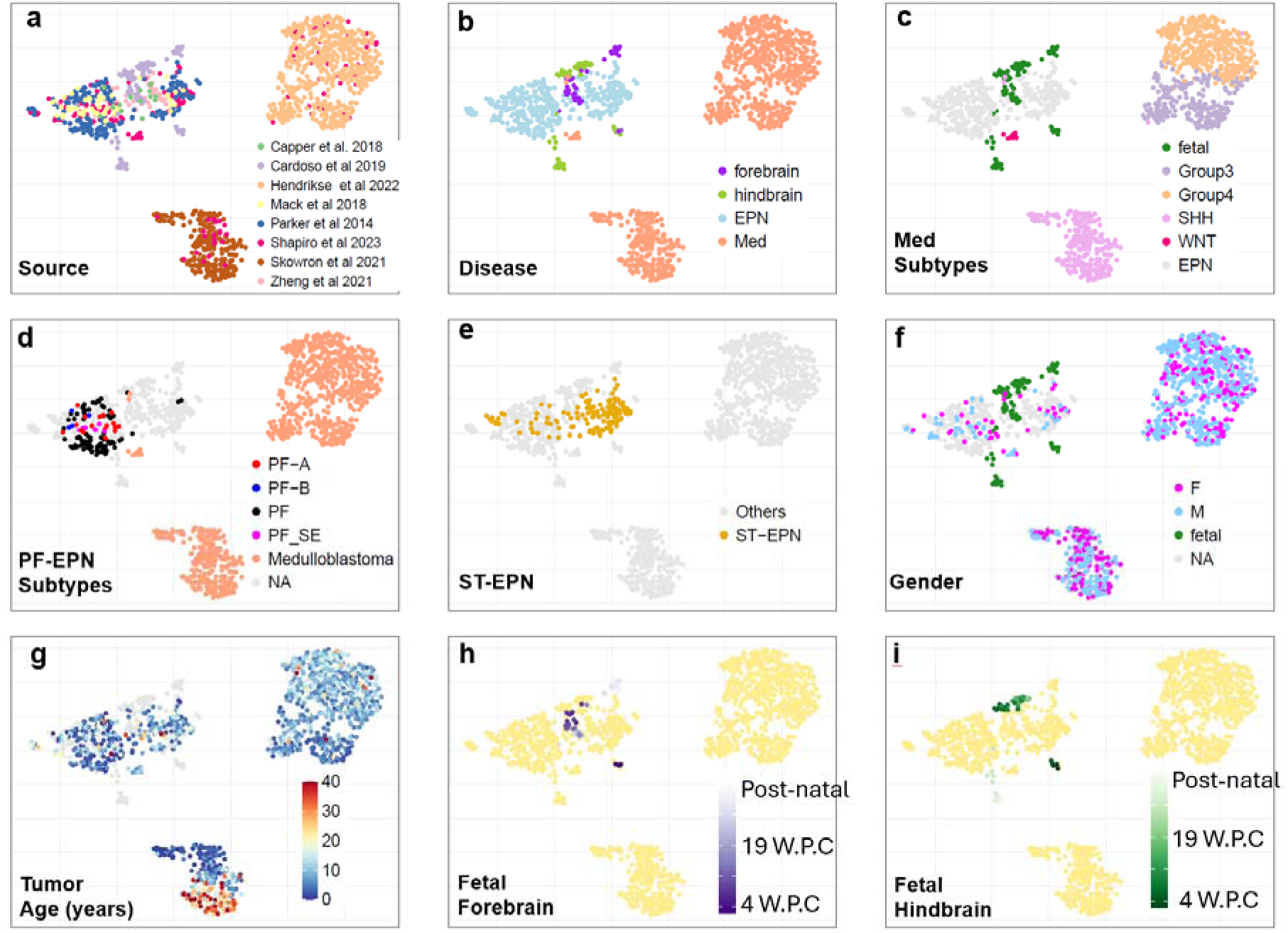
Generation of the Medulloblastoma and Ependymoma Landscape with Clinical and Genomic Metadata. (A) UMAP visualization colored by dataset source. (B) UMAP colored by disease type. (C) UMAP colored by subtypes for both medulloblastoma and ependymoma. (D) UMAP colored by subtypes within the posterior fossa. (E) UMAP highlighting supratentorial ependymomas (orange), with all other samples shown in grey. (F) UMAP colored by gender where available: Female (pink), Male (blue), and fetal samples (green). (G) UMAP colored by patient age at the time of tumor sample acquisition. (H) UMAP colored by age of forebrain samples. (I) UMAP colored by age of hindbrain samples.

### Overall structure of the reference landscape

Consistent with published analyses^2^, the medulloblastoma samples formed four distinct clusters corresponding to the SHH, Group 3, and Group 4 subtypes. Group 3 and Group 4 medulloblastomas were positioned along a continuum, consistent with previous reports. Notably, the nine WNT medulloblastoma samples clustered with the ependymoma samples, distinctly separate from the other medulloblastoma subtypes (**Fig. 1c**). The ependymoma clusters and the WNT medulloblastomas clustered closely with the developmental brain samples (**Fig. 1c**).

Coloring in the landscape based on the metadata collected for each sample, the ependymoma samples appeared to be divided into two major groups: posterior fossa ependymomas were predominantly localized in one cluster of the UMAP, while supratentorial ependymomas were primarily situated in the other (**Fig. 1d, e**). These two clusters corresponded to EPN-E1 and EPN-E2^19^. For samples where data was available, we further colored the posterior fossa ependymoma samples by annotated subtypes. The 17 PF-A samples were distributed across E2, the 4 PF-B samples formed a tight cluster at the top, and the 9 PF-SE (subependymoma) samples clustered in the middle of the posterior fossa region (**Fig. 1d**).

Overlaying sex information on our UMAP revealed no distinct regional patterns separating male and female samples on the reference landscape (**Fig. 1f**). However, when coloring the UMAP by the age of tumor samples, a clear age-based pattern emerged within the SHH medulloblastoma cluster. Specifically, older patients’ samples predominantly occupied one specific region of the cluster, while younger patients’ samples concentrated in the upper half, further validating our UMAP’s ability to differentiate between previously reported SHH medulloblastoma subgroups (**Fig. 1g**). Additionally, we visualized the age distribution for the forebrain and hindbrain samples (**Fig. 1h, i)** and noted that the early fetal forebrain and hindbrain samples clustered closely with the supratentorial ependymomas on the right side.

### Mapping Medulloblastoma Genomic Features onto the UMAP Landscape

While UMAP visualizations provide an intuitive representation of the data, these clusters are grounded in rigorous analyses performed in the original high-dimensional space and are further supported by biological validation using well-established genomic and transcriptomic markers from the literature. By Integrating known clinical features into the UMAP, distinct regionalization of different subtypes is revealed. We employed three methods: Arriba^20^ and STAR-Fusion^21^ to detect RNA fusions and CaSpER^22^ to infer copy number patterns for each sample in our dataset.

For medulloblastomas, it is known that the SHH (Sonic Hedgehog) subgroup is marked by significant deletions on chromosome 9q^7^, while Group 3 and Group 4 medulloblastomas are associated with a loss of chromosome 17p and a gain of 17q^9,23^. In our cohort, 32.12% (88/274) of SHH medulloblastomas exhibited a loss of chr9q (**Fig 2a**), 17.90% (41/229) of Group 3 medulloblastomas had a loss of chr17p, and 43.95% (160/364) of Group 4 medulloblastomas exhibited this loss (**Fig. 2b**). Furthermore, 62.08% (226/364) of Group 4 medulloblastomas and 22.70% (52/229) of Group 3 medulloblastomas had a gain of chr17q. (**Fig. 2c, Fig S1j**)

**Fig. 2.**
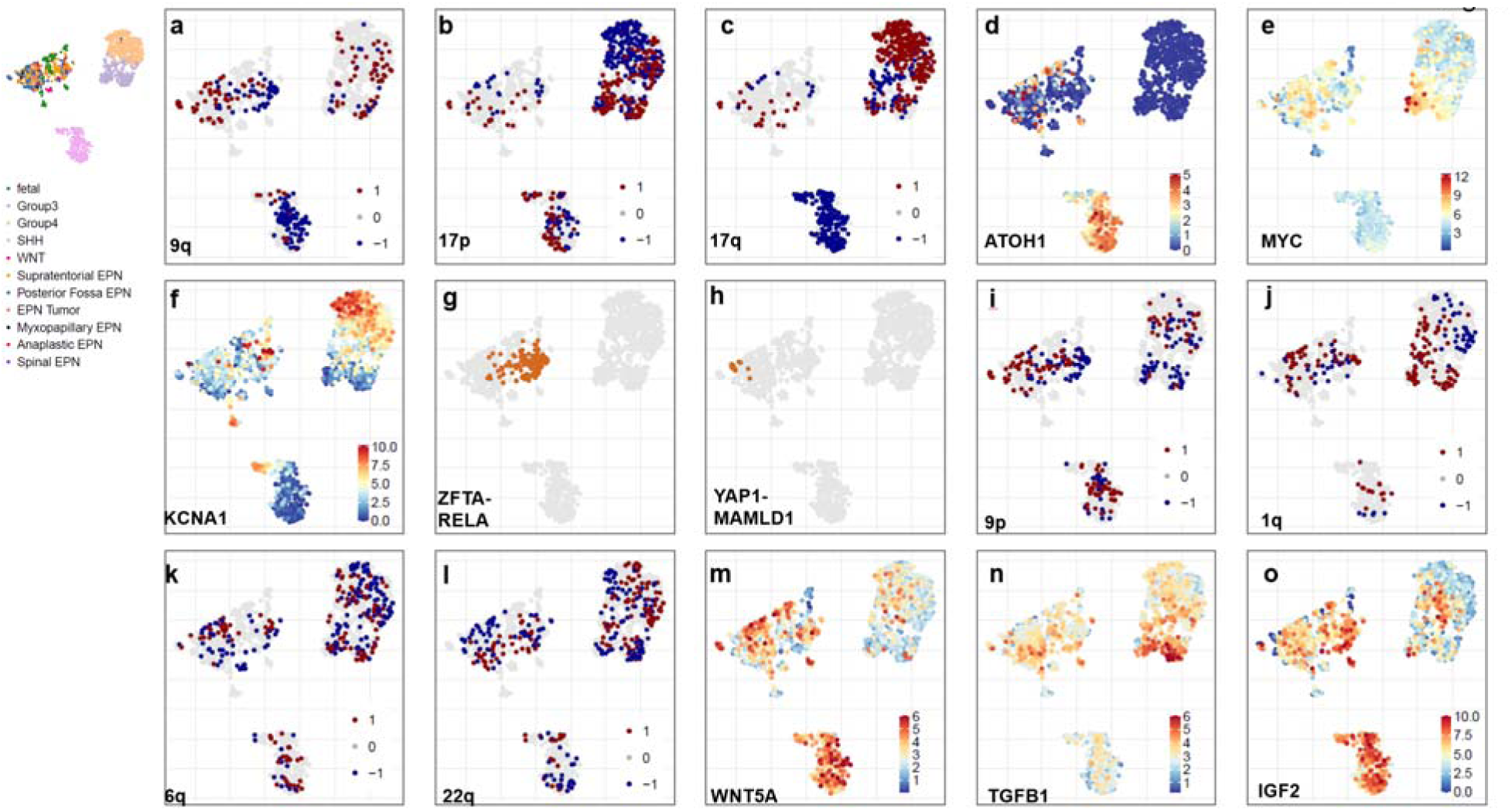
Validation of the reference landscape through copy number, gene fusions and gene expression patterns. (A-C) UMAP colored by copy number alterations to validate medulloblastoma subtypes: (A) 9q, (B) 17p, (C) 17q (red for gains, blue for deletions). (D-F) UMAP colored by gene expression levels: (D) ATOH1, (E) MYC, (F) KCNA1. (G-H) UMAP colored by gene fusions to confirm ependymoma subtypes: (G) ZFTA-RELA, (H) YAP-MAMLD1. (I-L) UMAP colored by copy number patterns in ependymomas: (I) 9p, (J) 1q, (K) 6q, (L) 22q. (M-O) UMAP colored by gene expression levels in ependymomas: (M) WNT5A, (N) TGFB1, (O) IGF2.

When we overlaid gene expression patterns onto the reference landscape, we observed distinct differences between SHH medulloblastomas and other tumor types. SHH medulloblastomas exhibited high expression of ATOH1 (**Fig. 2d, Fig S1k**), SFRP1 and HHIP (**Fig S2 a, b**), consistent with prior reports highlighting their elevated expression in this subtype^24^ In contrast, Group 3 medulloblastomas were characterized by elevated expression of MYC (**Fig. 2e**), GABRA5, and IMPG2 (**Fig S2 c, d**), while Group 4 medulloblastomas showed high expression of KCNA1 (**Fig. 2f**), EOMES, and RBM24 (**Fig S2. 2e,f**), corroborating existing literature that associates these genes with this subtype.

Consistent with previous studies, we identified several reported gene fusions across various medulloblastoma subtypes^24^. Specifically, as described by Luo et al, in our analysis 14.23% (39/274) of SHH medulloblastomas exhibited fusions involving CCDC196:LINC02290 **(Fig S2g)**. In contrast, among Group3 and Group4 medulloblastomas, 27% (62/229) of group 3 and 54% (200/364) of group4 showed gene fusion in GJE:VTA1, 6.9% (16/229) showed gene fusion in PVT1:PCAT1, 4.8%(11/229) of group 3 and 6.9% (25/364) of group4 showed gene fusion in TUBB2B:LMAN2L and 28/364(7.69%) of group4 in ELP4:IMMP1L (**Fig S2h-k**).

### Mapping Ependymoma Genomic Feature onto the UMAP Landscape

Supratentorial ependymomas (ST-EPNs) are frequently defined by specific gene fusions, most notably ZFTA-RELA^25^ and YAP1 fusions^5^, as well as recurrent losses of the entire chromosome 9 arm^5^. In our study, we observed that 71.64% (96/134) of ST-EPNs harbored the ZFTA-RELA fusion (**Fig. 2g**), while 6.71% (9/134) exhibited YAP1-MAMLD1 fusions. Notably, ZFTA-RELA fusions localized to a specific region on the UMAP, whereas YAP1-MAMLD1 fusions clustered on the opposite side, aligning more closely with posterior fossa ependymoma samples (**Fig 2h**). Additionally, 26.86% (36/134) of ST-EPN tumors showed deletions in the short arm of chromosome 9 (chr9p), and 23.88% (32/134) had deletions in the long arm (chr9q). (**Fig. 2a,i**).

By contrast, posterior fossa ependymomas are typically characterized by a gain of chromosome 1q^26^ and losses of chromosomes 6q and 22q^5^. Within our cohort, 12.59% (17/135) of posterior fossa ependymomas exhibited a 1q gain, 12.2% (17/134) had a 6q loss, and 11% (16/134) had a 22q loss (**Fig. 2j, k, l**). As previously reported, posterior fossa ependymomas exhibited elevated expression of WNT5A^26^ (**Fig. 2m**), TGFB1^26^ (**Fig2n**), and HOXB2^27^ (**Fig S2l.**), consistent with their established molecular signature. Similarly, supratentorial ependymomas demonstrated high expression of IGF2^28^ (**Fig. 2m**), L1CAM^29^, and CCND1^30^ (**Fig S2 n,o**), corroborating findings from earlier studies that identified these genes as key markers for this subtype. These results validate and extend existing knowledge of the gene expression profiles specific to ependymoma subtypes.

### Consensus clustering delineates stable molecular compartments in medulloblastoma

To define the transcriptional substructure of pediatric brain tumors, we performed consensus clustering on the full gene expression matrix, rather than the UMAP embedding. Unlike clustering in low-dimensional spaces like UMAP—which can distort distances and obscure subtle transcriptional differences—consensus clustering on the complete expression dataset provides a more stable and biologically meaningful stratification by assessing the reproducibility of sample groupings across multiple resampling iterations.

Using this approach, SHH medulloblastomas resolved into three distinct subclusters (**Fig. 3a**), aligning closely with previously described age-dependent subtypes. Group 3/4 medulloblastomas exhibited substantial heterogeneity, separating into six expression-based clusters (**Fig. 3b,c**), indicating a finer resolution of transcriptional diversity than previously reported.

**Fig. 3.**
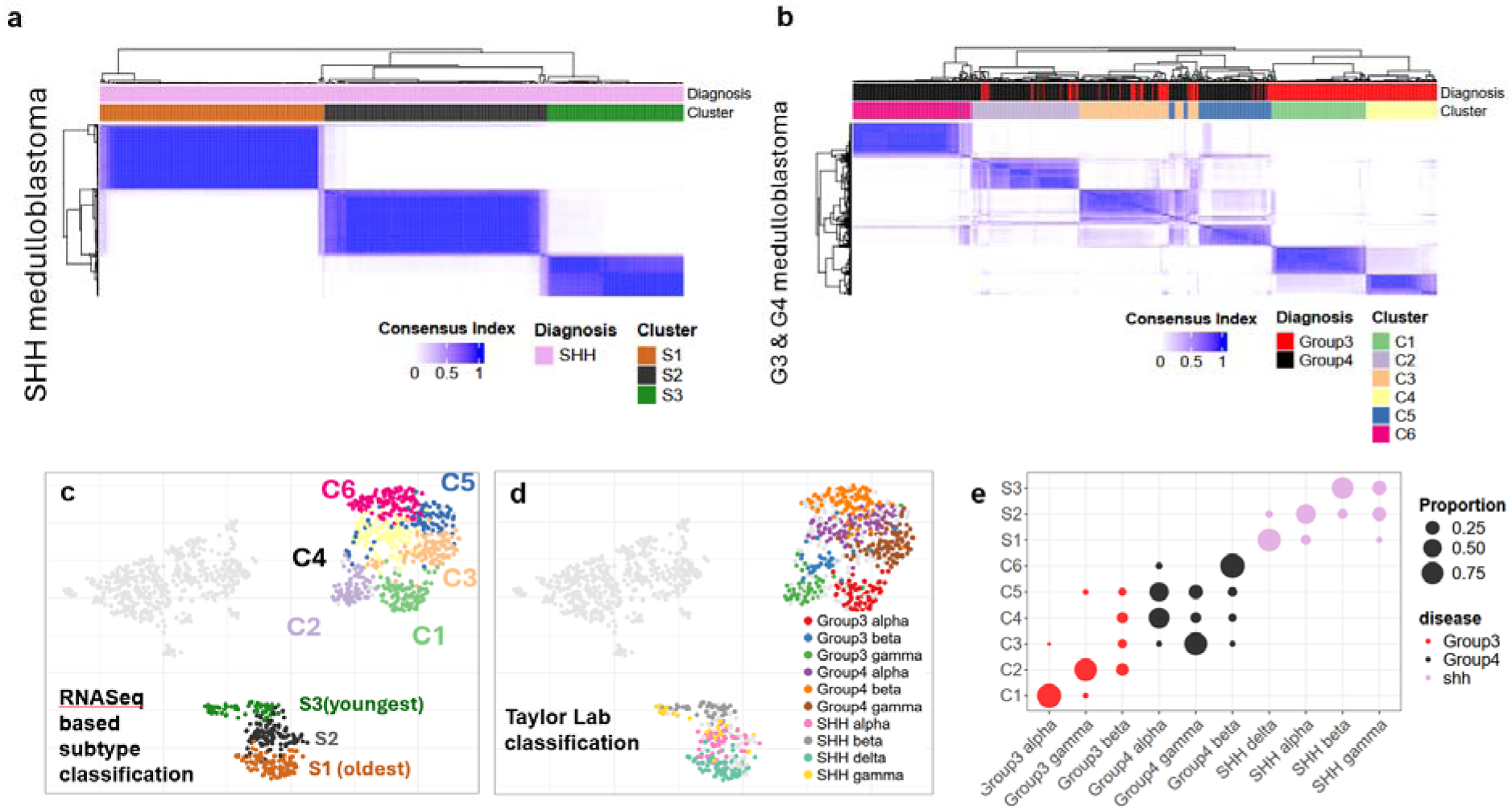
Consensus clustering reveals distinct subgroups of medulloblastoma. (A) Consensus clustering of shh medulloblastoma reveals three distinct clusters for SHH medulloblastoma, and (B) six different subclusters for group 3 and group4 medulloblastoma (D) UMAP colored by subtype as obtained from consensus clustering (D) UMAP colored by subtype classification according to Cavalli et al. (D) Dotplot showing the proportion of samples shared in Taylor Lab classification and RNASeq based consensus subtype classification

To benchmark our RNA-seq–based clustering, we compared it to the multi-omic classification described by Cavalli et al. (**Fig. 3d**). We found strong concordance between our clusters and the subtypes defined by Cavalli et al. and Skowron et al., with substantial sample overlap (**Fig. 3e**). Notably, our method—using only RNA-seq data—achieved comparable stratification of SHH medulloblastomas, accurately distinguishing S1 (SHHδ), S2 (SHHα), and S3 (SHHβ/γ) subgroups. These corresponded to median ages of 25, 6.3, and 1.6 years, respectively, highlighting the age-associated structure captured by our clustering.

In addition to transcriptional resolution, our approach revealed clinically relevant associations. Patients in S2 with 9q deletions exhibited significantly improved survival compared to copy-neutral cases (p = 0.0029, **Fig. S3a**). As previously reported by Cavalli et al., S2 also showed enrichment for 9p amplifications, 9q deletions, and 10q deletions, while S1 exhibited a higher frequency of 14q deletions (Fig. S3b, Table S2). These genomic features reinforce the robustness of our RNA-based subtyping.

Group 3/4 tumors were similarly stratified into six distinct clusters. Group 3 tumors split into C1, C2, and C4, while Group 4 tumors segregated into C3, C5, and C6. Consistent with Cavalli et al., C2 showed strong MYC upregulation and frequent chromosome 8 gains (8p: 52.77%, 8q: 63.88%; Fig. 2e, **Fig. S3c**, Table S2). By contrast, C1 was marked by gain of chromosome 7 (7p: 25%, 7q: 37.5%), loss of chromosome 8 (8p: 39.58%, 8q: 31.25%), and gain of 14q (52.08%) (Table S2). Subgroup C3 exhibited widespread loss of both 8p (68.46%) and 8q (63.06%) (Table S2).

Further, we also noted a subgroup of C4 demonstrated significant sex-based survival differences, with males faring worse than females (p = 0.015, **Fig S3d**). Additionally, C4 patients with deletions in chromosome 4p had better outcomes than those who were copy-neutral for this region (p = 0.032, **Fig. S3e**, Table S2). Isochromosome 17 also displayed varied copy number patterns across clusters: 17q gain was seen in C1 (21.87%), C2 (23.6%), C3 (47.47%), C4 (46.36%), C5 (52.94%), and C6 (76.47%), while 17p loss was most prominent in C1 (16.6%) (Table S2).

### Bulk and Single-Cell RNA-Seq Concordantly Define Subtype-Specific Expression patterns and Pathways in Medulloblastoma

To uncover biologically distinct signaling programs within medulloblastoma, we gene set variation analysis (GSVA) on RNA-seq data from each molecular subtype. This revealed unique transcriptional signatures and signaling pathway enrichments distinguishing SHH, Group 3, and Group 4 tumors.

Across all SHH tumors, we identified distinct pathways that were uniquely upregulated in this subtype, distinguishing them from other medulloblastoma subtypes. Specifically, pathways involving Gli protein binding to promoters, RUNX3 regulation of YAP1-mediated transcription, ribosome-related processes, and B-lymphocyte signaling were prominently upregulated in SHH tumors. We also noted additional upregulated pathways, such as protein kinase C activity, metabolic processes, wound healing, nonsense mediated decay, and T-cell activation (**Fig. 3a-f, Fig S3f**).

The Group 4 subgroups (C3, C5, and C6) displayed distinct pathway regulation compared to the Group 3 subgroups. Specifically, the MYC-amplified Group 3 subgroup C2 was enriched for pathways related to translation, Wnt signaling, phototransduction activation, TERT pathway, and voltage-gated channels (**Fig. 3f-h, Fig S4a-e**). In contrast, Group 4 subgroups showed upregulation in pathways such as NTRK2 signaling, STAT5 activation, signaling by leptin, IL22BP pathway, KIT signaling, and presynaptic depolarization and calcium signaling (**Fig. 3i-l, Fig S4h-i**).

To validate the novel pathways identified in each subtype, we analyzed single-cell RNA sequencing (scRNA-seq) data from 25 patients, comprising 5 WNT, 3 SHH, 8 Group 3, and 9 Group 4 samples, obtained from Hovestadt et al. Following data preprocessing using Seurat, we observed that WNT and SHH subtypes formed distinct clusters, whereas Group 3 and Group 4 samples exhibited closer clustering patterns (**Fig. S4j**). This spatial relationship mirrors the clustering observed in the UMAP projection generated from bulk RNA-seq data. Differential gene expression analysis further corroborated subtype-specific patterns consistent across both scRNA-seq and bulk RNA-seq datasets. For instance, SHH samples exhibited elevated expression of ATOH1, SFRP1, and HHIP, while Group 3 displayed overexpression of MYC, GABRA5, and IMPG2, and Group 4 demonstrated higher expression of KCNA1, EOMES, and RBM24 (**Fig. S4k,l**). Gene set variation analysis (GSVA) reinforced the activation of novel pathways identified in scRNA-seq data, aligning with findings from bulk RNA-seq. Notably, NTRK2 signaling was most prominent in Group 4, and **WNT** signaling was enriched in Group 3, among other pathway-specific enrichments (**Fig. S4m**).

**Fig. 4.**
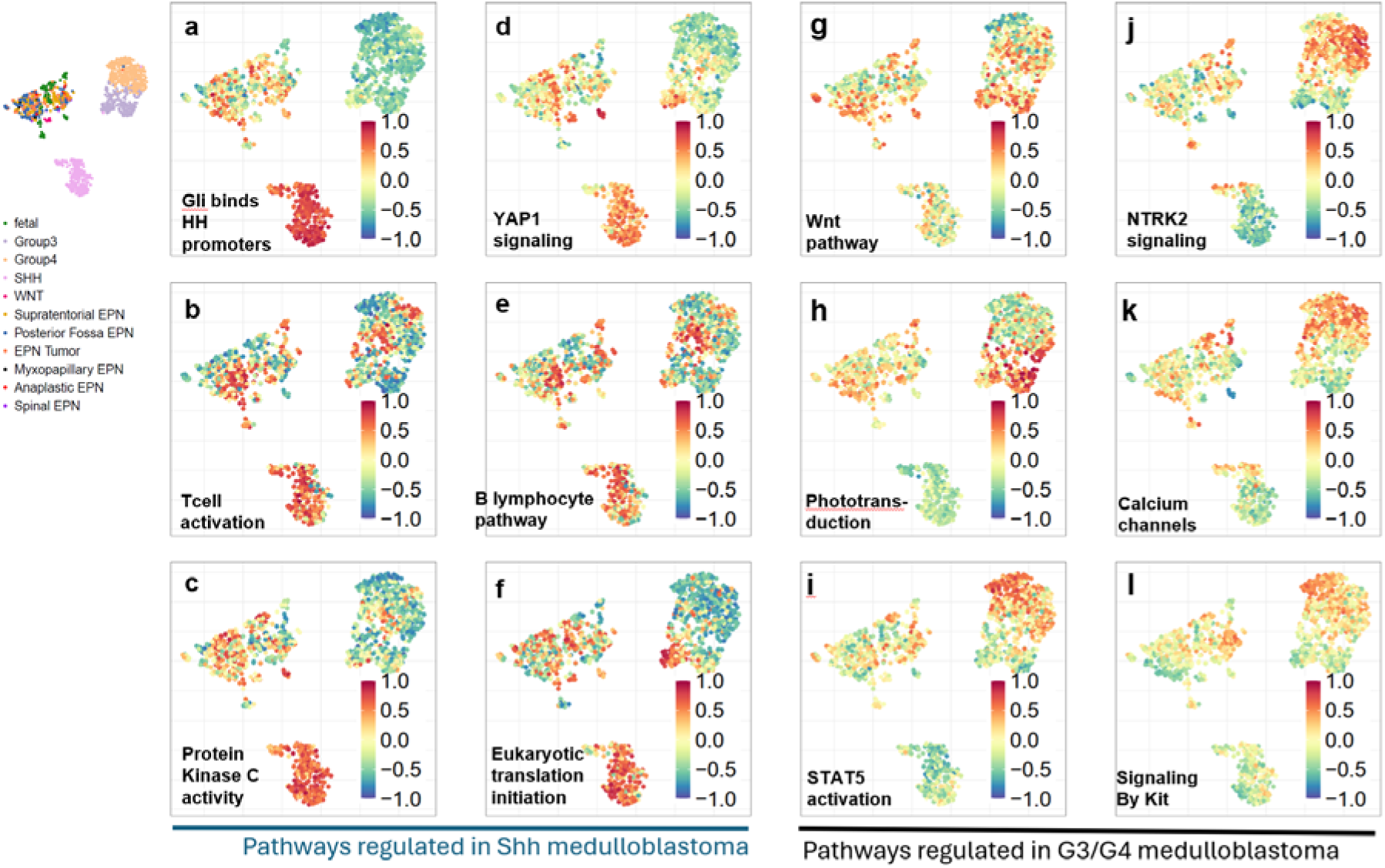
Group 3 and Group 4 Medulloblastoma Subtypes with Distinct Molecular Profiles Identified by RNA-Seq. (A-F) Pathways upregulated in SHH medulloblastoma compared to Group 3 and Group 4 medulloblastoma samples(G-L). UMAP colored in by GSVA score, A score closer to 1 denotes up-regulation of pathway and closer to –1 denotes down-regulation of pathway.

### Clustering Reveals Two Distinct Ependymoma Subgroups with distinct gene fusions

To validate and build upon prior molecular subtype classifications, we reanalyzed the same ependymoma samples previously studied by Chan et al^19^. who identified two robust consensus clusters—EPN-E1 and EPN-E2—using bulk RNA-seq data.

Reapplying consensus clustering to this dataset, we recapitulated this two-subgroup structure (**Fig. 5a)**. To ensure subtype-specific resolution, WNT medulloblastoma and fetal brain samples were excluded from the ependymoma clustering analysis. When visualized using UMAP, the samples colored by EPN-E1 and EPN-E2 assignments formed two clearly separable clusters, confirming the reproducibility of these transcriptional subtypes in our dimensionality-reduced reference landscape (**Fig. 5b)**. To rule out the possibility that the observed transcriptomic differences could be due to variations in tumor purity rather than genuine biological differences, we estimated tumor purity and cell composition using the PUREE^31^ *and* ESTIMATE^32^ methods. Neither method showed differences across clusters in tumor purity or stromal infiltration (**Fig S5a,b**). We observed that the forebrain and hindbrain samples clustered closely to the ependymoma samples, we thus performed a correlation analysis of EPN samples using top8000 most variable genes and observed a strong correlation with early developmental time points, as opposed to later stages (**Fig S5c, d**).

**Fig. 5.**
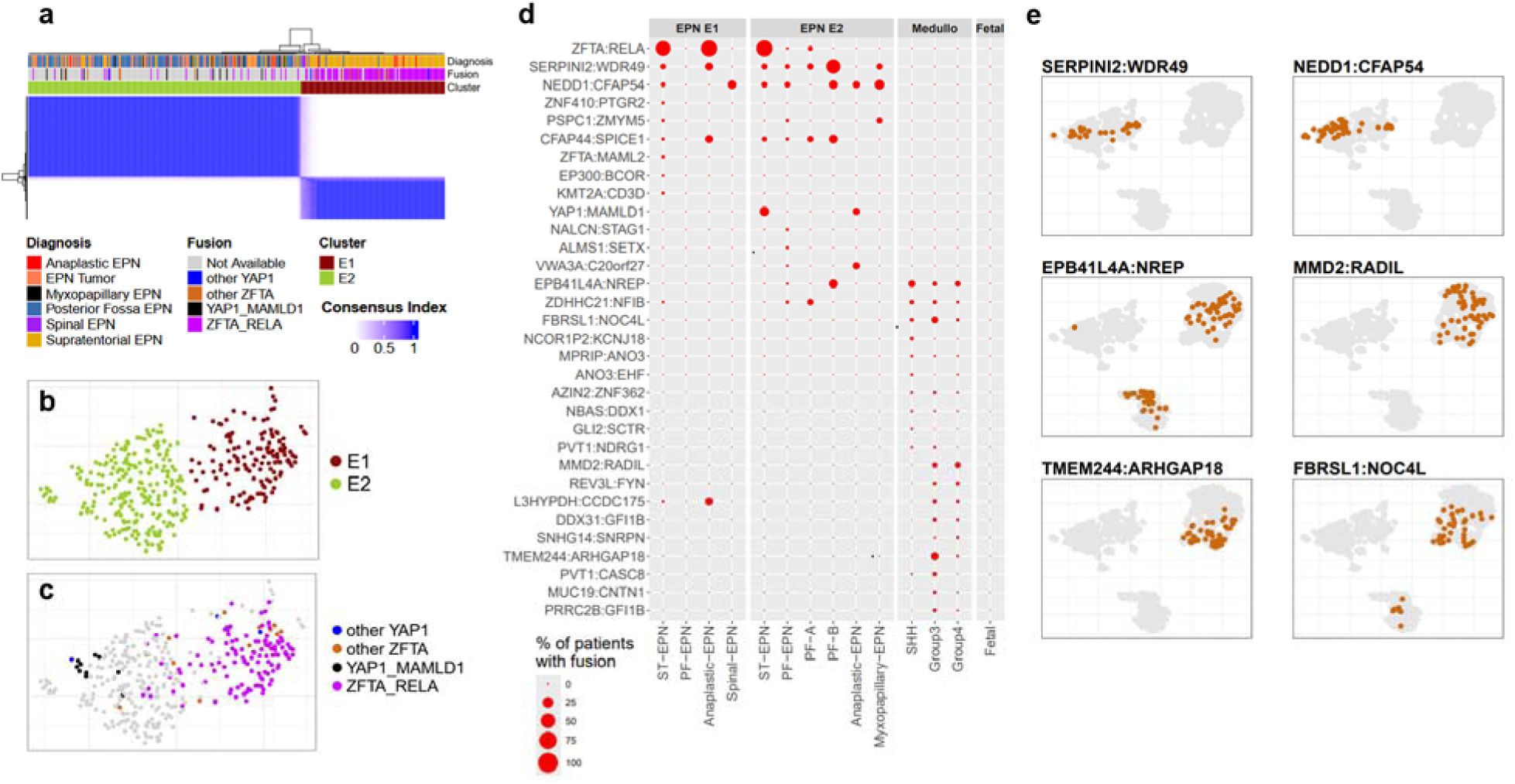
Ependymoma segregate into two clusters EPN-E1 and EPN-E2. (A) Consensus clustering of ependymoma reveals two clusters, EPN-E1 and EPN-E2. (B UMAP colored by subtype EPN-E1 and EPN-E2 as obtained from consensus clustering. (C) UMAP showing the distribution of commonly studied gene fusions in ependymoma: ZFTA-RELA (purple), YAP1-MAMLD1 (black), other YAP1 fusions (blue), and other ZFTA fusions (brown). (D) Dot plots showing the gene fusions and their frequencies in EPN-E1 and EPN-E2, as well as across medulloblastoma subtypes. (E) UMAP colored in by patients displaying gene fusions regionalized in distinct ependymoma and medulloblastoma subtypes.

EPN-E1 was predominantly composed of supratentorial ependymomas (ST-EPNs), comprising 77% of the cluster, whereas EPN-E2 was enriched for posterior fossa ependymomas (PF-EPNs, 55%), ST-EPNs (12%), ependymoma NOS samples (25%), myxopapillary EPNs (3.85%) and Anaplastic EPNs (2.56%). Tumors diagnosed as myxopapillary ependymomas are exclusively localized within or near the EPN-E2 cluster, whereas spinal and anaplastic ependymomas were distributed across both EPN-E1 and EPN-E2 (**Fig. 5b**). We note that while some of the diagnosis reported here may be outdated with respect to the 2021 WHO classification, we have presented the metadata as originally annotated to ensure transparency and consistency.

A subset of ependymomas have recurrent chromosomal translations that generate oncogenic gene fusions^13^, which have been shown to drive tumorigenesis in mouse models^25^. We analyzed gene fusion profiles across EPN-E1 and EPN-E2 clusters and observed distinct patterns between them. The EPN-E1 cluster, predominantly composed of ST-EPNs, was enriched for the canonical ZFTA:RELA gene fusion, found in 82% of samples (87/105; **Table S3a**). Among EPN-E2, 31.0% (9/29) harbored a YAP1:MAMLD1 fusion, and 27.6% (8/29) carried a ZFTA:RELA fusion (**Fig. 5c, Table S3b**). This suggests that while ZFTA:RELA remains a dominant marker in EPN-E1, a portion of ST-EPNs in EPN-E2 also harbor this fusion, indicating that gene fusion status alone is not uniquely causal for generation of the EPN-E1 gene expression pattern.

Further investigation of the ST-EPNs with ZFTA:RELA fusions in EPN-E2 revealed no other recurrent gene fusions (**Table S3c**). All five anaplastic ependymomas in EPN-E1 carried the ZFTA:RELA fusion (**Table S3d**), whereas the six anaplastic cases in EPN-E2 showed no ZFTA:RELA or other recurrent fusions (**Table S3e**).

As previously described, we also identified non-RELA ZFTA fusions in EPN-E1, including ZFTA:NCOA2 (3/105, 2.9%) and ZFTA:MAML2 (3/105, 2.9%) (Fig. 5d). By contrast, EPN-E2 showed distinct fusions not typically associated with ST-EPNs, such as NEDD1:CFAP54, observed in 10.1% (13/129) of PF-EPNs (**Table S4a, Fig. 5e**). The small number of PF-EPNs present in EPN-E1 (n=5) did not exhibit any recurrent gene fusions (Table S4b). Due to limited sample sizes, we were unable to identify fusion trends in PF-A (n=17), PF-B (n=4), or myxopapillary ependymomas (n=9) (**Tables S4c–e**).

By contrast, the medulloblastoma samples exhibited a higher frequency of RNA fusions per sample compared to ependymomas (**Fig S5h**) and differed markedly from that of the ependymomas, potentially generated by DNA rearrangement gene fusion or RNA trans-splicing mechanisms. While EPBH41L4A:NREP ( 14%, 5.2% and 8.79% in SHH, Group3 and group4 medulloblastoma respectively, **Table S5a-c**) was found in all three medulloblastoma subtypes, MMD2:RADIL ( 7.42% and 10.16% respectively), TMEM244:ARGHAP18(19.65% and 0.82% respectively) and FBRSL1:NOC4L (14.41% and 2.74% respectively) fusions were seen only in grade 3 and grade 4 subtypes. (**Fig5e**).

### Distinct Gene and Pathway Regulation in EPN-E1 and EPN-E2 subgroups

Differential gene expression analysis between EPN-E1 and EPN-E2 revealed 106 kinases were upregulated in EPN-E1 and another 105 kinases were up-regulated in EPN-E2 (**Fig. 6a**, highlighted in pink). These included tyrosine receptor kinases, such as oncogenic driver MERTK and EPHB4 which were up-regulated in EPN-E1 (**Fig 6b**) and (**Table S6a**) and NTRK2/3 (**Fig 6b**) up-regulated in EPN-E2, which play an oncogenic role in adult glioma^33^ and several other cancer types^34^. Of note, MerTK was found to be an oncogenic driver in the ZFTA-RELA fusion-driven mouse model of EPN^19^. The gene expression profiles of E2 ZFTA-RELA tumors are significantly different from the E1 ZFTA-RELA tumors **(Fig S6a).** Notably, the E1 non-ZFTA-RELA tumors show greater similarity to E1 ZFTA-RELA tumors than to E2 ZFTA-RELA tumors **(Fig S6b)**.

**Fig. 6.**
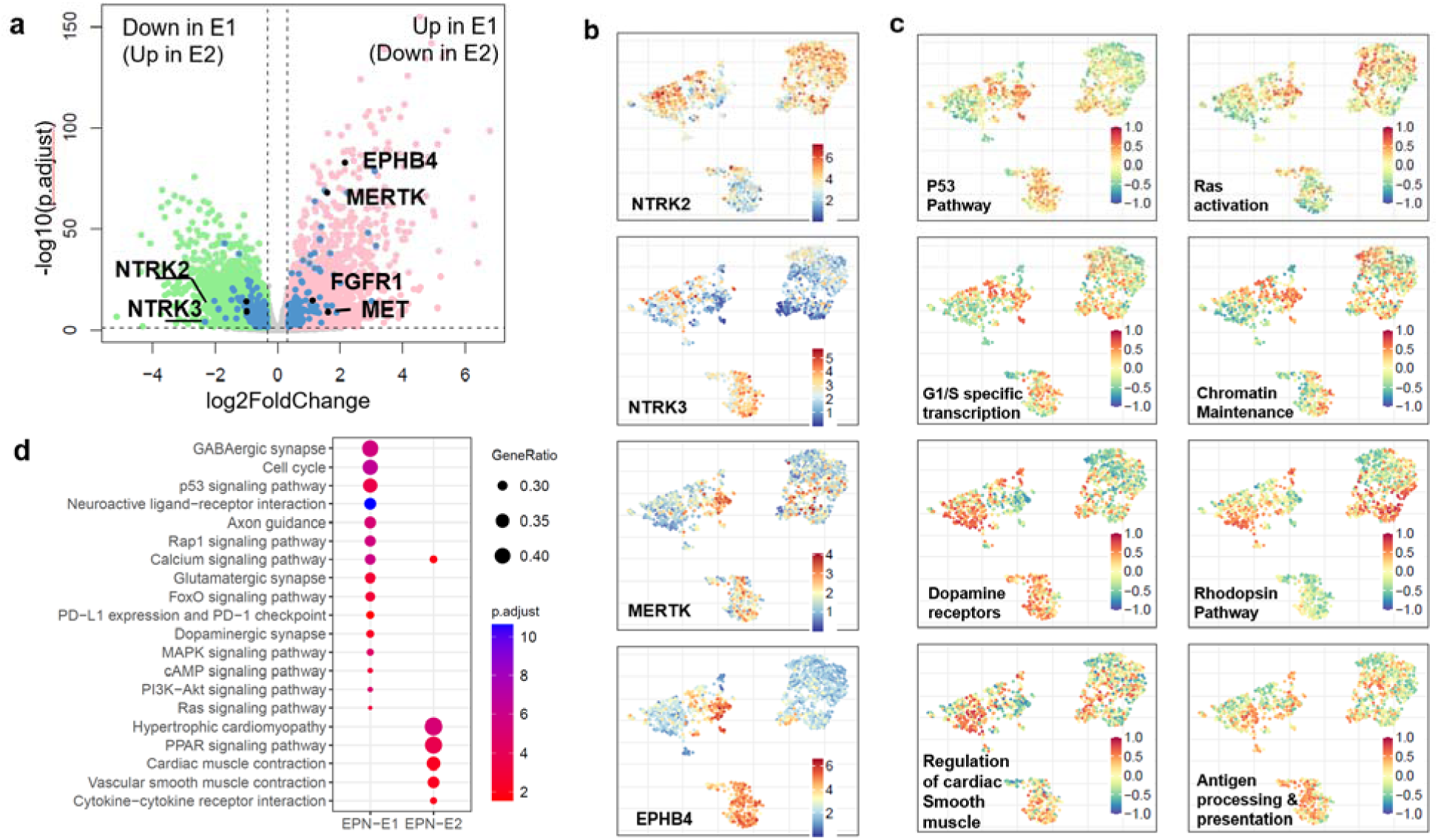
Contrasting Differences Between EPN-E1 and EPN-E2. (A) Volcano plots showing differentially expressed genes in EPN-E1 (pink) and EPN-E2 (green), with differentially regulated kinases highlighted in blue, the top tyrosine kinase receptors are labeled in black. (B) Gene expression levels of tyrosine receptor kinases: NTRK2, NTRK3, MERTK, and EPHB4. (C) UMAP showing GSVA scores for pathways regulated in EPN-E1. (E) UMAP showing GSVA scores for pathways regulated in EPN-E2. (D) Dot plot illustrating pathways upregulated in EPN-E1 and EPN-E2.

Several synaptic genes which are critical for neuronal communication, neurodevelopment and cognitive development^35^ were also differentially up-regulated in EPN-E1 compared to EPN-E2, such as GRIN1, CHRNB1, CACNA1G, CACN1B and P2RX5 (**Fig S6c-g)**. EPN-E2 also had a distinct set of up-regulated synaptic markers such as GABRA5, DRD1, SCN4B and P2RX7 **(Fig S6h-k)**.

Further analysis revealed distinct pathway regulation between the EPN-E1 and EPN-E2 groups. EPN-E1 showed upregulation of pathways involved in Notch signaling, the TP53 pathway, RAS signaling, and interferon gamma (IFNG) signaling (**Fig. 6c,d, Table S7a**). Additionally, pathways related to chromatin maintenance and G1/S-specific transcription were upregulated in EPN-E1. By contrast, EPN-E2 exhibited upregulation of pathways associated with hyaluronan biosynthesis, dopamine receptor signaling, voluntary skeletal muscle contraction, and antigen processing and presentation (**Fig. 6c,d, Table S7b**). Single-cell RNA sequencing studies^36,37^ have revealed that ependymoma tumor samples contain subpopulations with ependymal cell characteristics and transitions into mesenchymal-like subpopulations. Therefore, we did a GSVA analysis to determine if there was a pattern of mesenchymal character in either tumor cluster that would explain the expression separation between the two. While some gene sets were more expressed in one compared to the other, there was no overall pattern of mesenchymal expression that would explain the separation of the two (**Fig S6l**).

### Projecting New Patients onto a Pre-existing UMAP Reference Landscape

Our established UMAP landscape has clearly delineated distinct biological regions corresponding to different disease types and subtypes. This landscape can also be leveraged to overlay new patients entering the clinic, aiding in the prediction of their disease subtype and ruling out misdiagnosis. To demonstrate this concept, we utilized previously developed algorithm^18^ to project new patient data onto the reference landscape.

One of the primary data sources for our landscape is the Children’s Brain Tumor Network (CBTN), which includes 77 ependymoma and 93 medulloblastoma samples among 23 different pediatric tumor types. Of the 93 medulloblastoma samples, 10 were classified as Group 3, 38 as Group 4, 24 as SHH, 9 as WNT, and 12 as NOS. We employed a nearest neighbors algorithm to overlay 12 NOS samples (A-L; Table **S8**, **Fig. 7a**) onto our landscape. The results showed that patients H and I co-embedded within the EPN-E2 group, while patient F clustered with the WNT medulloblastomas. Patients A and G aligned with the MYC-amplified Group C2, and patient L corresponded with C1. Patient C localized to C4, whereas patients B and D were clearly situated within Group 4 (C6 and C5). Additionally, patients K, E, and J were positioned at the boundary between Group 3 and Group 4, within C5 and C3.

**Fig. 7.**
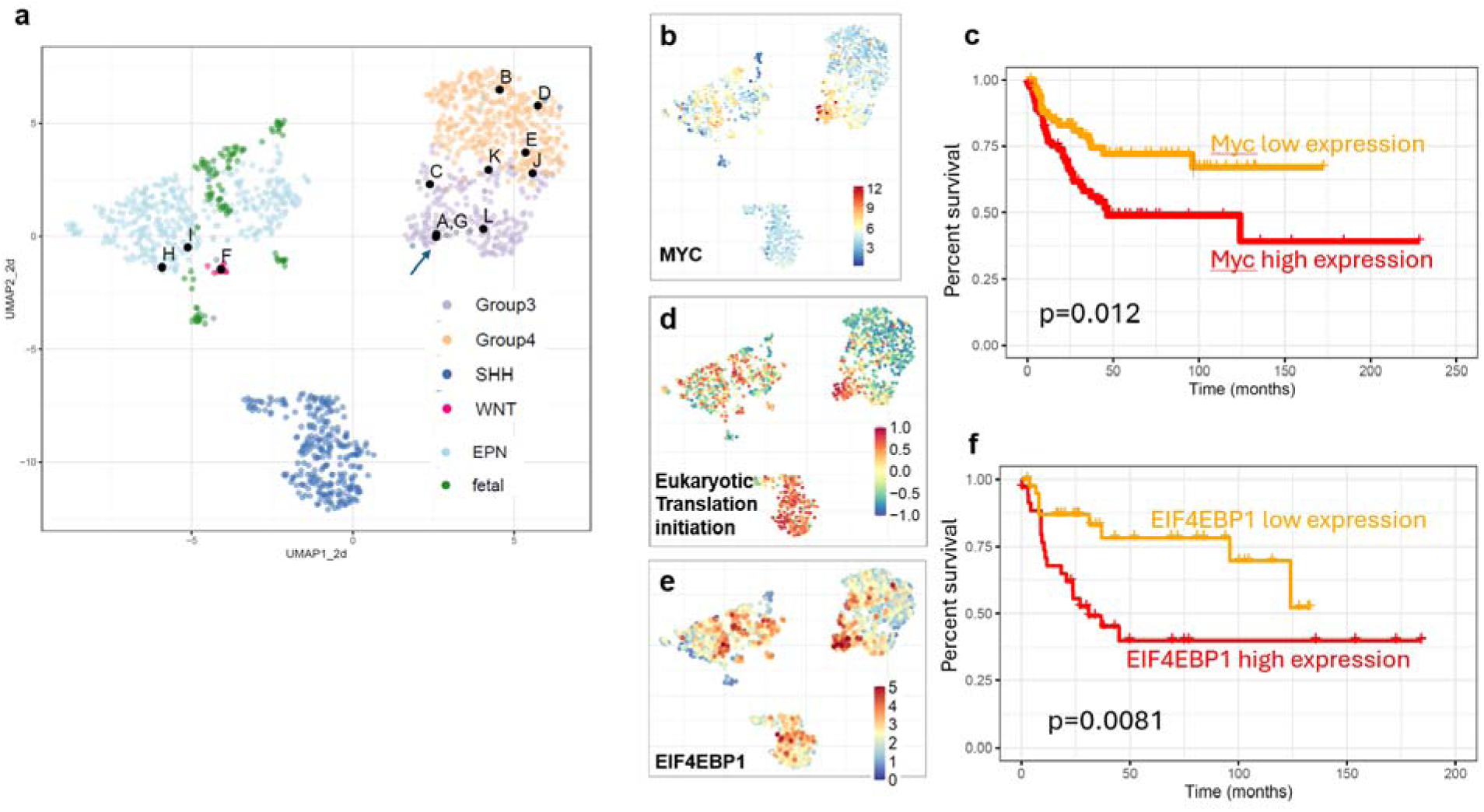
Integrating New Patient Data onto the Reference Landscape. (A) Using the k-nearest neighbors algorithm to assign subtypes to 12 NOS medulloblastoma samples. (B) UMAP colored by MYC gene expression. (C) Kaplan Meier plots for C2 based on MYC gene expression levels (High(red) vs Low(orange), pvalue = 0.012, n[high exp]=35, n[low exp]=109). (D) UMAP displaying GSVA scores for eukaryotic translation pathways. (E) Gene expression levels of EIF4EBP1 on the UMAP. (F) Kaplan Meier plots for C2 based on EIF4EBP1 gene expression levels (High(red) vs Low,(orange) pvalue = 0.0081, n[high exp]=14, n[low exp]=77).

To validate the accuracy of our algorithm, we examined the molecular profiles of the 6 out of 12 NOS tumors that overlapped with the Group 3clusters. For instance, the median MYC expression value in C2 is 7.5 (log2(TPM)). Patients A, G, J, and L, all of whom fell within the MYC-amplified region, exhibited MYC expression levels of 6.56, 7.33, 7.28, and 6.25, (log2(TPM)) respectively (**Fig. S7**). Additionally, patient A showed a gain of chromosome 8q, consistent with the profile of Group 3 subtypes. Patients C and K, located at the boundary between Group 3 and Group 4 tumors, also displayed elevated MYC expression (6.21 and 4.9, respectively), with patient K showing a gain of 8q.

Similarly, the Group 4 subclusters C6 and C5 are characterized by high EOMES expression, with a median expression value of 6.17 and 5.31 (log2(TPM)) respectively. Evaluating the 3 out of 12 NOS Patients B, D, and E that overlapped with Group4 cluster, we observed that they exhibited elevated EOMES expression (6.37, 4.10, and 6.4, (log2(TPM)) respectively). Patient B also demonstrated a gain of 17q, aligning with the Group 4 subtype profile. We recalculated the UMAP with the 12 medulloblastoma NOS samples included and found that the predicted placement based on the nearest neighbors algorithm (shown in black) precisely matched the ground truth (red) in terms of their positioning on the landscape. (**Fig S7**)

Overlaying new patient data onto the landscape can also inform therapeutic decisions. For example, within the C2 group, characterized by high MYC expression, we found that stratifying patients based on MYC levels revealed poor prognosis in patients with elevated MYC expression levels, as reported previously^38^(p = 0.012, **Fig 7b,c**). This subgroup also exhibited upregulation of pathways involved in translation (**Fig. 7d**). EIF4EBP1 is a known negative regulator of translation initiation, and its elevated levels have been linked to drug resistance^39^. Notably, EIF4EBP1 was similarly overexpressed in C2, mirroring the MYC expression pattern. When we further stratified patients based on EIF4EBP1 expression, those with high levels of EIF4EBP1 had poorer survival compared to those with lower expression (p = 0.081, **Fig 7e,f**). Therefore, if a new patient, such as patient A (**Fig. 7**), falls into a region associated with high EIF4EBP1 expression, they may be at increased risk for drug resistance. Importantly, our landscape helps identify co-occurring transcriptional programs such as Myc high expression and translation.

## DISCUSSION

Over the past decade, the emergence of single-cell atlases^40,41^. and bulk RNASeq derived reference landscapes^10,18^ aimed at molecularly characterizing various diseases has become increasingly prevalent. Though landscapes derived from single-cell RNA sequencing (scRNA-seq) reveals cellular heterogeneity, it demands significant resources, time, and complex analysis.

By contrast, landscapes developed from publicly available bulk RNA-seq datasets offer a cost-effective and efficient alternative for building reference landscapes. Leveraging publicly available data enables rapid construction of robust landscapes, facilitating exploration of molecular targets and disease subtypes, and accelerating research and discovery.

The landscape approach to studying a disease has significant advantages. Firstly, the comprehensive nature of our landscape, built from a large number of samples across various tumor types, uncovers novel tumor biology. Through our analysis, we have uncovered previously unrecognized gene fusion events in specific tumor subtypes, pathway regulation that contribute to our understanding of disease mechanisms and could serve as new therapeutic targets. For example, the up-regulation of translation in SHH, STAT5 and NTRK2 signaling in Group4 and up-regulation of TERT pathway and WNT pathway in Group3, underscores the potential of these landscapes in refining disease classification and improving our understanding of tumor heterogeneity.

Secondly, our study identifies distinct clustering patterns, such as the EPN-E1 and EPN-E2 clusters in ependymomas. The 2 clusters, validated by multiple clustering approaches show distinct gene fusions and pathway regulation. These patterns help delineate subtypes of diseases, offering valuable insights into the underlying biology. For example, we observed distinct synaptic genes and tyrosine receptor kinases upregulated in each of the clusters. While several tyrosine receptor kinases are now actionable targets in precision oncology, synaptic genes are also increasingly being viewed as therapeutic targets. Drugs that modulate synaptic function, such as those targeting neurotransmitter receptors or synaptic proteins may offer treatments for diseases. These findings suggest the presence of distinct tumor-neuron synaptic signaling pathways in EPN-E1 and EPN-E2, which have not yet been explored in ependymomas.

Thirdly, these landscapes offer significant clinical value, particularly in the context of constructing clinical trials. By overlaying new patient data onto the reference landscape, clinicians can assess a patient’s molecular profile based on their nearest neighbors within the landscape, helping to refine and infer diagnoses. This approach is especially useful when traditional diagnostics are inconclusive or in cases of atypical disease presentations. Additionally, integrating new patient data into these landscapes can help identify potential misdiagnoses, ensuring that patients receive the most accurate and effective treatment based on their specific disease subtype.

Given that medulloblastoma is heavily studied, it served as a valuable control, allowing us to validate our approach by confirming well-documented findings from the literature. Additionally, by analyzing medulloblastoma and ependymoma together, we identified not only disease-specific alterations but also unexpected transcriptional similarities between certain subtypes of the two entities. While distinct gene fusions were observed in each tumor type, we found that several genes and pathways were regulated in a strikingly similar manner across subtypes. For example, CHRNA4, a neuronal nicotinic acetylcholine receptor, mutations in which cause epilepsy, was upregulated in EPN-E1 and SHH medulloblastoma, whereas GABRA5, a GABA-A receptor, responsible for mediating inhibitory neurotransmission, was upregulated in EPN-E2 and Group 3 (MYC-amplified medulloblastoma). These shared transcriptional patterns suggest potential biological parallels, providing new perspectives on pediatric CNS tumors. This comparative approach not only enhances our understanding of tumor biology but could also highlight potential shared therapeutic targets across these diseases.

Lastly, by making the landscape available as a freely accessible online tool, researchers are provided with a toolbox to explore genes of interest within the context of our reference landscape, facilitating the discovery of new biomarkers and enhancing the broader scientific community’s ability to conduct hypothesis-driven research. By providing this resource, we aim to empower researchers with the tools necessary to uncover novel insights into disease biology, ultimately contributing to the development of more effective treatments.

In conclusion, our study highlights the value of bulk RNA-seq-derived reference landscapes as a cost-effective and powerful tool for disease characterization, diagnostic refinement, and the identification of novel molecular features. As the field continues to evolve, integrating such landscapes into both research and clinical settings will be crucial in advancing our understanding and treatment of complex diseases.

## MATERIALS AND METHODS

### Collection of publicly available RNA Sequencing data

Raw RNA sequencing data for medulloblastoma and ependymoma samples were retrieved from various public data repositories, as detailed in Table S1. The Heidelberg dataset was obtained from the data repository of the Department of Neuropathology at the University Hospital Heidelberg.

### RNA-Seq data processing and visualization

Quality assessment of the raw RNA sequencing data was performed using FastQC (v0.11.9) in conjunction with MultiQC (v1.9) to generate comprehensive reports. The RNA sequencing reads were then aligned to the Gencode GRCh38.primary_assembly reference genome using STAR^42^ (v2.7.7a). Gene-level quantification was conducted with HTSeq^43^ (v0.11.0) using Gencode^44^ V39 primary assembly annotations. The raw gene counts from all datasets were subsequently aggregated and batch effects were corrected using the ComBat-seq function from the R package “sva”^45^. Normalized gene expression values were calculated and expressed as VST from “DESeq2”^46^ package.

### Clustering Approaches

Dimensionality reduction via Uniform Manifold Approximation and Projection (UMAP) was applied to the VST normalized expression data from protein-coding genes to construct the medulloblastoma and ependymoma reference landscape. UMAPs were generated using the “umap” R package (https://cran.r-project.org/web/packages/umap/index.html). The following parameters were used for the umap function: n_neighbors=15, n_components=2 (for 2d umap) and n_components = 3 (for 3d umap present on Oncoscape), metric = Euclidean, n_epichs = 200 and min.disct = 0.5). consensusClusteringPlus^47^ was used to determine the optimum k for shh medulloblastoma, Group3/group4 medulloblastoma and ependymoma samples. WNT medulloblastoma and fetal samples were removed when sub clustering the ependymoma samples.

### Gene Fusion Detection from RNA-Seq

Gene fusions were identified using Arriba^20^ (v2.1.0) and STAR-Fusion^21^ on STAR two-pass aligned RNA-Seq data. Only high-confidence fusions from both tools and involving at least one protein-coding gene (per GENCODE v39, GRCh38.p14) were retained.

### Copy Number Alterations (CNA) Detection from RNA-Seq

Large-scale copy number alterations, including chromosome arm-level changes, were inferred for all tumors using the CaSpER^22^ package applied to bulk RNA-Seq data. BAFExtract, including its source code, genome list, and genome pileup directory, was obtained from https://github.com/akdess/. Cytoband and centromere data for the hg38 reference genome were sourced from the UCSC Genome Browser.

### Kaplan-Meier Survival Analysis

Kaplan-Meier survival curves were generated using the recurrence data for each sample, focusing only on tumors with known recurrence status and known time to recurrence or last follow-up. Kaplan-Meier curves were plotted, and p-values were calculated using the R package “survival” (v3.5.7).

### Differential gene expression analysis

Differential expression analysis was performed using DESeq2^46^. Significantly regulated genes in each comparison were identified based on FDR (< 0.05) and log 2fold change (> 0.3) or fold change of 25%

### GSVA Pathway Analysis

Pathway gene sets from KEGG^48^, Biocarta^49^, Reactome^50^ pathways and Gene Ontology Biological Processes were sourced from the Molecular Signatures Database (MSigDB) version 7.2 (https://www.gsea-msigdb.org/gsea/msigdb/collections.jsp). Gene Set Variation Analysis (GSVA)^51^ was performed on batch-corrected VST counts for all samples. The resulting GSVA scores, ranging from 1 to –1 for each sample, were visualized using ggplot2^52^.

### Placing new patients on UMAP reference map

VST counts for the 12 medulloblastoma NOS samples were calculated and used as test data. The VST counts for all remaining samples were used as training data. The k-nearest neighbors algorithm, developed in Thirimane et al^18^, which overlays new patient data onto an existing UMAP based on its nearest neighbors was used to predict the location of the test samples on the reference umap. The obtained UMAP coordinates were then added to the existing UMAP object and plotted using ggplot2 in R.

### Oncoscape integration

Matrix and clinical data were prepared for Oncoscape by converting them to cBioPortal formats (cbioportal.org). Custom settings, including colorings and precalculated views to match the paper’s figures, were stored in JSON in an Oncoscape updates.txt file. See https://github.com/FredHutch/OncoscapeV3/blob/master/docs/upload.md for details.

## Acknowledgements

We thank members of the Holland lab at Fred Hutch Cancer Center for valuable discussions and collaborators for sharing their data and metadata. This research was supported by funding from the Fred Hutch Cancer Center (E.C.H), 1R35 CA253119-01A1 (E.C.H).

## Funding

National Institutes of Health grant 1R35 CA253119-01A1 (E.C.H).

## Author Contributions

Conceptualization, S.A., and E.C.H.; Methodology, S.A., and E.C.H.; Formal Analysis, S.A. and N.N.; Software, M.J., G,G; Investigation, S.A., E.C.H.; Resources, M.D.T.; Data Curation, S.A. and N.N., Writing – Original Draft, S.A and E.C.H; Writing – Review & Editing, S.A., E.C.H; Visualization, S.A., M.J., G,G; Supervision E.C.H., Funding Acquisition, E.C.H.

## Declaration of interests

Although the majority of Oncoscape has been open source for many years, a provisional patent has been filed on subset of the technology and computational algorithms presented in this paper, and N.N, S.A, M.J and E.C.H are listed as inventors (Serial No.: 63/595,717).

## Data and materials availability

All analysis including statistics and visualization were done in R version 4.3. Plots were generated using R basic graphics and ggplot2. Raw sequencing data was downloaded from E-MTAB-6814, GSE109381, EGAS00001002696, EGAS00001000254, EGAD00001006305 and EGAS00001005826. All custom code used in this study are available at https://github.com/sonali-bioc/MedulloEPNLandscape

**Table S1.** Datasets from North America and Europe were combined to generate the medulloblastoma UMAP.

**Table S2.** Copy number profiles for each subtype of Group3 and Group4 medulloblastoma, and each subtype of shh medulloblastoma, as obtained from consensus clustering.

**Table S3.** Top recurrent RNA gene fusions in ST-EPNs in EPN-E1.

**Table S4.** Top recurrent RNA gene fusions in PF-EPNs in EPN-E2.

**Table S5.** Top recurrent RNA fusions in medulloblastoma subtypes.

**Table S6.** All genes and kinases upregulated in EPN-E1 and EPN=E2.

**Table S7.** Pathways upregulated in EPN-E1 and EPN-E2.

**Table S8.** Mapping of 12 NOS medulloblastoma samples from CBTN as shown in Figure 7.

## Notes

https://github.com/sonali-bioc/MedulloEPNLandscape

### Competing Interest Statement

The authors have declared no competing interest.

### Summary of Updates

Update to figures, new analysis added

## References and Notes

1. Brown, N.J., et al. The 100 Most Influential Publications on Medulloblastoma: Areas of Past, Current, and Future Focus. World Neurosurg 146, 119–139 (2021).

2. Taylor, M.D., et al. Molecular subgroups of medulloblastoma: the current consensus. Acta Neuropathol 123, 465–472 (2012).

3. Pajtler, K.W., et al. YAP1 subgroup supratentorial ependymoma requires TEAD and nuclear factor I-mediated transcriptional programmes for tumorigenesis. Nat Commun 10, 3914 (2019).

4. Mack, S.C., et al. Therapeutic targeting of ependymoma as informed by oncogenic enhancer profiling. Nature 553, 101–105 (2018).

5. Pajtler, K.W., et al. Molecular Classification of Ependymal Tumors across All CNS Compartments, Histopathological Grades, and Age Groups. Cancer Cell 27, 728–743 (2015).

6. Vladoiu, M.C., et al. Childhood cerebellar tumours mirror conserved fetal transcriptional programs. Nature 572, 67–73 (2019).

7. Skowron, P., et al. The transcriptional landscape of Shh medulloblastoma. Nat Commun 12, 1749 (2021).

8. Suzuki, H., et al. Recurrent noncoding U1 snRNA mutations drive cryptic splicing in SHH medulloblastoma. Nature 574, 707–711 (2019).

9. Cavalli, F.M.G., et al. Intertumoral Heterogeneity within Medulloblastoma Subgroups. Cancer Cell 31, 737–754 e736 (2017).

10. Arora, S., et al. Visualizing genomic characteristics across an RNA-Seq based reference landscape of normal and neoplastic brain. Sci Rep 13, 4228 (2023).

11. McFerrin, L.G., et al. Analysis and visualization of linked molecular and clinical cancer data by using Oncoscape. Nat Genet 50, 1203–1204 (2018).

12. Capper, D., et al. DNA methylation-based classification of central nervous system tumours. Nature 555, 469–474 (2018).

13. Parker, M., et al. C11orf95-RELA fusions drive oncogenic NF-κB signalling in ependymoma. Nature 506, 451–455 (2014).

14. Shapiro, J.A., et al. OpenPBTA: An Open Pediatric Brain Tumor Atlas. bioRxiv (2022).

15. Zheng, T., et al. Cross-Species Genomics Reveals Oncogenic Dependencies in ZFTA/C11orf95 Fusion-Positive Supratentorial Ependymomas. Cancer Discov 11, 2230–2247 (2021).

16. Hendrikse, L.D., et al. Failure of human rhombic lip differentiation underlies medulloblastoma formation. Nature 609, 1021–1028 (2022).

17. Cardoso-Moreira, M., et al. Gene expression across mammalian organ development. Nature 571, 505–509 (2019).

18. Thirimanne, H.N., et al. Meningioma transcriptomic landscape demonstrates novel subtypes with regional associated biology and patient outcome. Cell Genom 4, 100566 (2024).

19. Chan, M., et al. A Systems approach identifies MERTK as a therapeutic vulnerability in ZFTA-RELA-driven ependymomas. bioRxiv, 2025.2002.2006.636949 (2025).

20. Uhrig, S., et al. Accurate and efficient detection of gene fusions from RNA sequencing data. Genome Res 31, 448–460 (2021).

21. Haas, B.J., et al. Accuracy assessment of fusion transcript detection via read-mapping and de novo fusion transcript assembly-based methods. Genome Biol 20, 213 (2019).

22. Serin Harmanci, A., Harmanci, A.O. & Zhou, X. CaSpER identifies and visualizes CNV events by integrative analysis of single-cell or bulk RNA-sequencing data. Nat Commun 11, 89 (2020).

23. Northcott, P.A., et al. The whole-genome landscape of medulloblastoma subtypes. Nature 547, 311–317 (2017).

24. Luo, Z., et al. Genomic and Transcriptomic Analyses Reveals ZNF124 as a Critical Regulator in Highly Aggressive Medulloblastomas. Front Cell Dev Biol 9, 634056 (2021).

25. Ozawa, T., et al. A De Novo Mouse Model of C11orf95-RELA Fusion-Driven Ependymoma Identifies Driver Functions in Addition to NF-κB. Cell Rep 23, 3787–3797 (2018).

26. Gödicke, S., et al. Clinically relevant molecular hallmarks of PFA ependymomas display intratumoral heterogeneity and correlate with tumor morphology. Acta Neuropathol 147, 23 (2024).

27. Zaytseva, M., Papusha, L., Novichkova, G. & Druy, A. Molecular Stratification of Childhood Ependymomas as a Basis for Personalized Diagnostics and Treatment. Cancers (Basel) 13(2021).

28. Korshunov, A., et al. Gene expression patterns in ependymomas correlate with tumor location, grade, and patient age. Am J Pathol 163, 1721–1727 (2003).

29. Chavali, P., et al. L1CAM Immunopositivity in Anaplastic Supratentorial Ependymomas: Correlation With Clinical and Histological Parameters. Int J Surg Pathol 27, 251–258 (2019).

30. Torre, M., et al. Characterization of molecular signatures of supratentorial ependymomas. Mod Pathol 33, 47–56 (2020).

31. Revkov, E., Kulshrestha, T., Sung, K.W. & Skanderup, A.J. PUREE: accurate pan-cancer tumor purity estimation from gene expression data. Commun Biol 6, 394 (2023).

32. Yoshihara, K., et al. Inferring tumour purity and stromal and immune cell admixture from expression data. Nat Commun 4, 2612 (2013).

33. Pattwell, S.S., et al. A kinase-deficient NTRK2 splice variant predominates in glioma and amplifies several oncogenic signaling pathways. Nat Commun 11, 2977 (2020).

34. Pattwell, S.S., et al. Oncogenic role of a developmentally regulated NTRK2 splice variant. Sci Adv 8, eabo6789 (2022).

35. Michetti, C., Falace, A., Benfenati, F. & Fassio, A. Synaptic genes and neurodevelopmental disorders: From molecular mechanisms to developmental strategies of behavioral testing. Neurobiol Dis 173, 105856 (2022).

36. Gillen, A.E., et al. Single-Cell RNA Sequencing of Childhood Ependymoma Reveals Neoplastic Cell Subpopulations That Impact Molecular Classification and Etiology. Cell Rep 32, 108023 (2020).

37. Gojo, J., et al. Single-Cell RNA-Seq Reveals Cellular Hierarchies and Impaired Developmental Trajectories in Pediatric Ependymoma. Cancer Cell 38, 44–59.e49 (2020).

38. Grotzer, M.A., et al. MYC messenger RNA expression predicts survival outcome in childhood primitive neuroectodermal tumor/medulloblastoma. Clin Cancer Res 7, 2425–2433 (2001).

39. Schuster, S.L. & Hsieh, A.C. The Untranslated Regions of mRNAs in Cancer. Trends Cancer 5, 245–262 (2019).

40. Cao, J., et al. The single-cell transcriptional landscape of mammalian organogenesis. Nature 566, 496–502 (2019).

41. Domcke, S., et al. A human cell atlas of fetal chromatin accessibility. Science 370(2020).

42. Dobin, A., et al. STAR: ultrafast universal RNA-seq aligner. Bioinformatics 29, 15–21 (2013).

43. Anders, S., Pyl, P.T. & Huber, W. HTSeq--a Python framework to work with high-throughput sequencing data. Bioinformatics 31, 166–169 (2015).

44. Frankish, A., et al. GENCODE reference annotation for the human and mouse genomes. Nucleic Acids Res 47, D766–d773 (2019).

45. Leek, J.T., Johnson, W.E., Parker, H.S., Jaffe, A.E. & Storey, J.D. The sva package for removing batch effects and other unwanted variation in high-throughput experiments. Bioinformatics 28, 882–883 (2012).

46. Love, M.I., Huber, W. & Anders, S. Moderated estimation of fold change and dispersion for RNA-seq data with DESeq2. Genome Biol 15, 550 (2014).

47. Wilkerson, M.D. & Hayes, D.N. ConsensusClusterPlus: a class discovery tool with confidence assessments and item tracking. Bioinformatics 26, 1572–1573 (2010).

48. Kanehisa, M. & Goto, S. KEGG: kyoto encyclopedia of genes and genomes. Nucleic Acids Res 28, 27–30 (2000).

49. BioCarta. Biotech Software & Internet Report 2, 117–120 (2001).

50. Gillespie, M., et al. The reactome pathway knowledgebase 2022. Nucleic Acids Res 50, D687–d692 (2022).

51. Sonja Hänzelmann, R.C.J.G. GSVA: gene set variation analysis for microarray and RNA-Seq data. BMC Bioinformatics (2013).

52. Wickham, H. ggplot2: Elegant Graphics for Data Analysis, (Springer-Verlag New York, 2016).

