## Supplementary material for "Integrated transcriptomic landscape of medulloblastoma and ependymoma reveals novel tumor subtype-specific biology": Supp figs

Supplementary Information

Table of contents

Supplementary Figures S1-S7

Supplementary Figure Legends

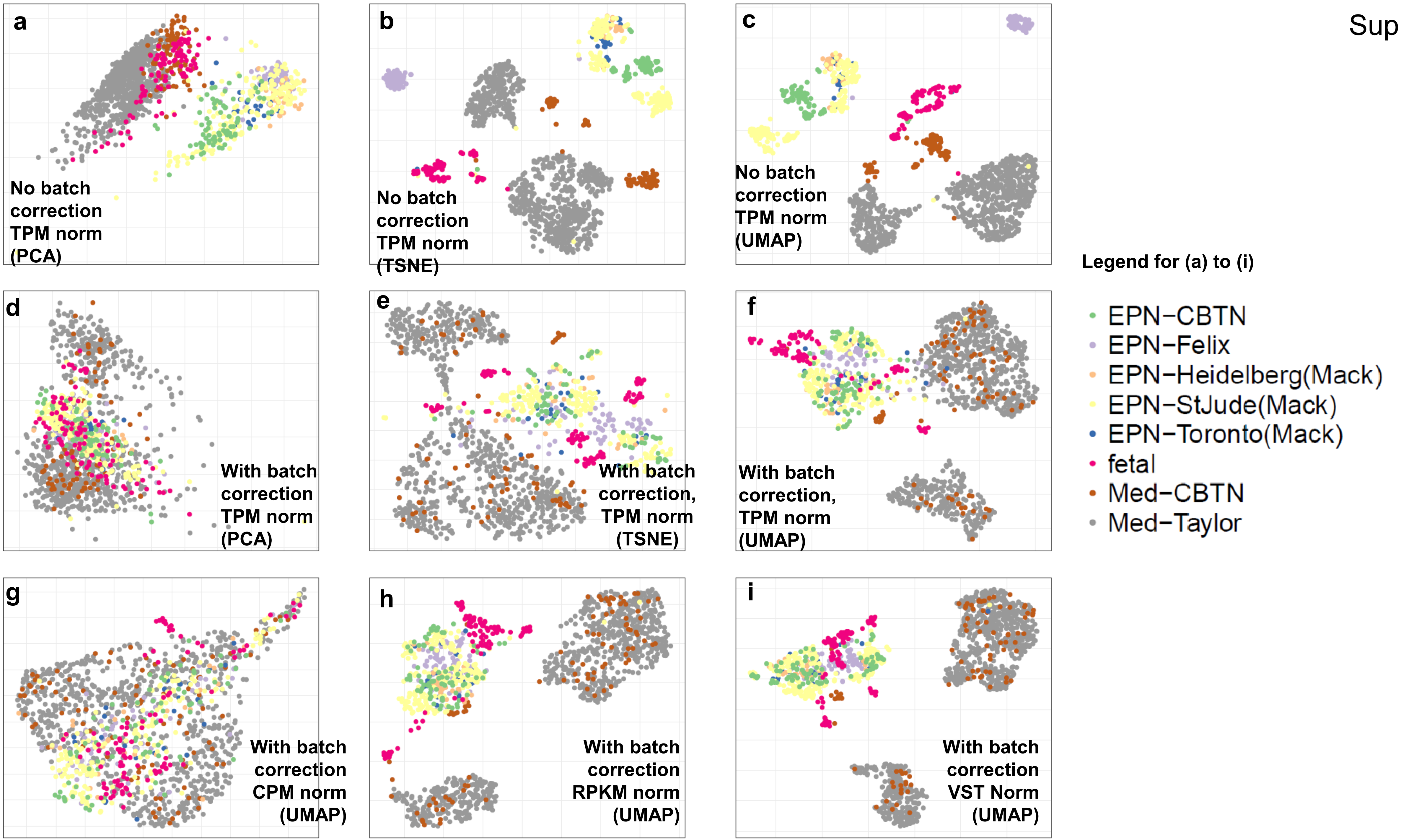
a
No batch correction TPM norm (PCA)
b
No batch correction TPM norm (TSNE)
c
No batch correction TPM norm (UMAP)
d
With batch correction TPM norm (PCA)
e
With batch correction, TPM norm (TSNE)
f
With batch correction, TPM norm (UMAP)
g
With batch correction CPM norm (UMAP)
h
With batch correction RPKM norm (UMAP)
i
With batch correction VST Norm (UMAP)
Legend for (a) to (i)
EPN−CBTN
EPN−Felix
EPN−Heidelberg(Mack)
EPN−StJude(Mack)
EPN−Toronto(Mack)
fetal
Med−CBTN
Med−Taylor

j

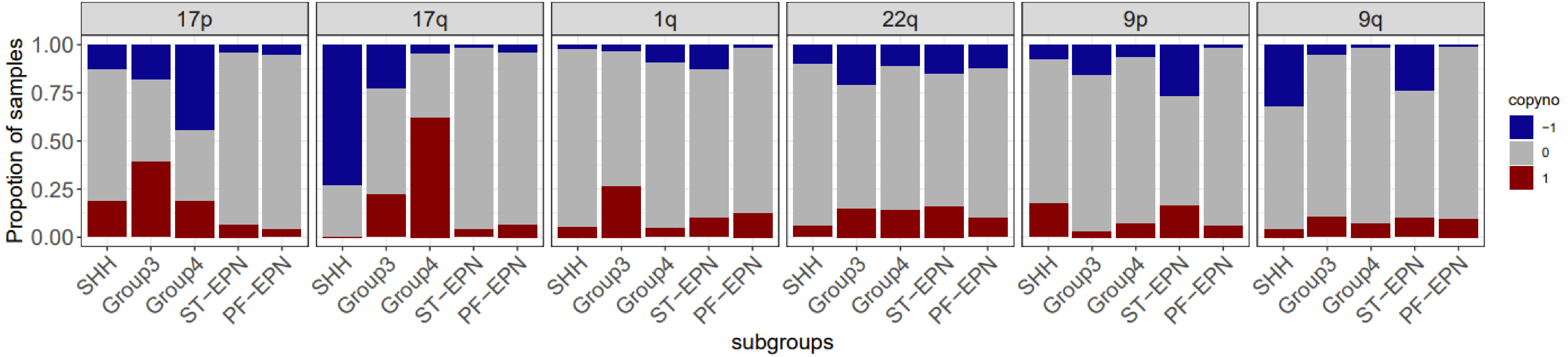
17p
17q
1q
22q
9p
9q
Propotion of samples
1.00
0.75
0.50
0.25
0.00
SHH
Group3
Group4
ST–EPN
PF–EPN
subgroups
copyno
–1
0
1

k

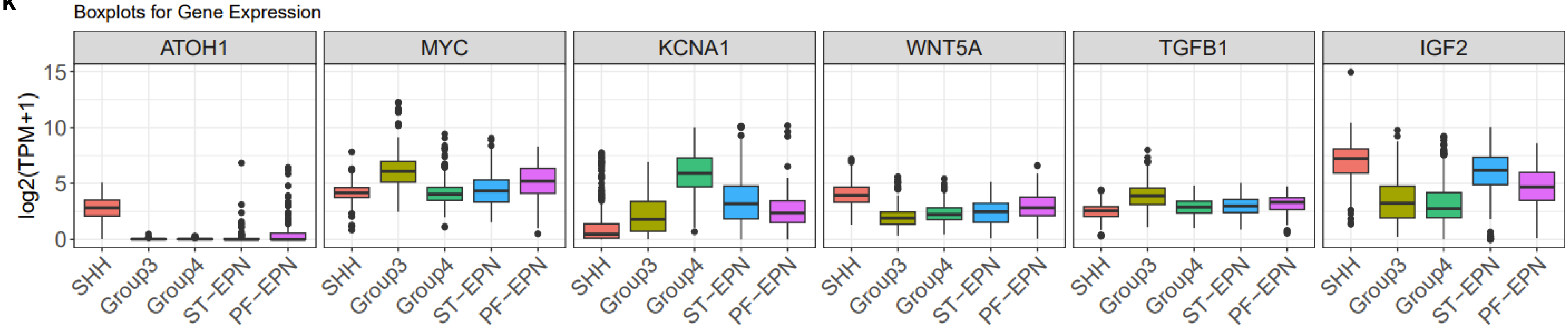
Boxplots for Gene Expression
ATOH1
MYC
KCNA1
WNT5A
TGFB1
IGF2
log2(TPM+1)
15
10
5
0
SHH
Group3
Group4
ST–EPN
PF–EPN

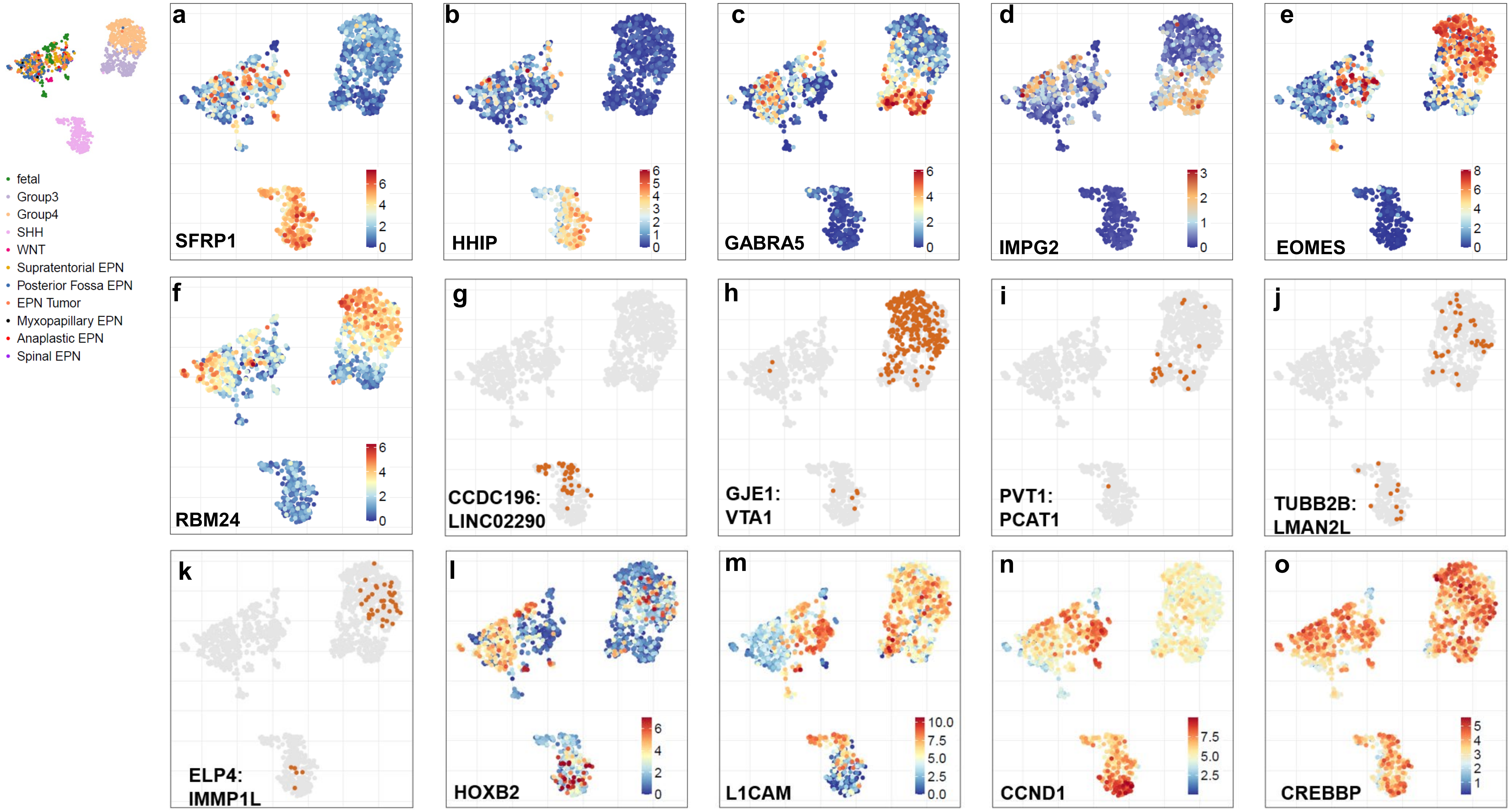
fetal
Group3
Group4
SHH
WNT
Supratentorial EPN
Posterior Fossa EPN
EPN Tumor
Myxopapillary EPN
Anaplastic EPN
Spinal EPN
a
SFRP1
b
HHIP
c
GABRA5
d
IMPG2
e
EOMES
f
RBM24
g
CCDC196:
LINC02290
h
GJE1:
VTA1
i
PVT1:
PCAT1
j
TUBB2B:
LMAN2L
k
ELP4:
IMMP1L
l
HOXB2
m
L1CAM
n
CCND1
o
CREBBP

Supp Fig 3

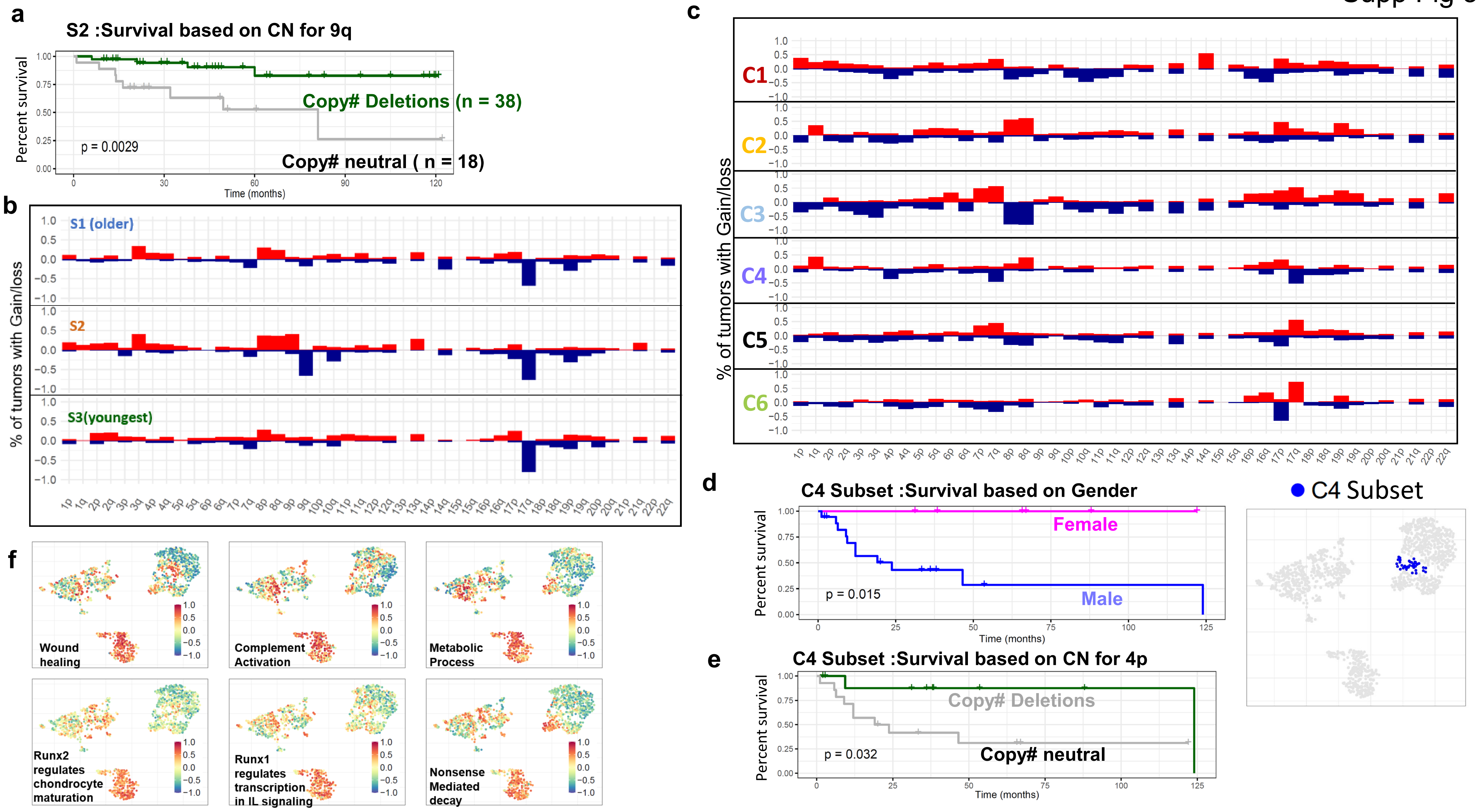
a
S2 :Survival based on CN for 9q
Percent survival
Copy# Deletions (n = 38)
p = 0.0029
Copy# neutral ( n = 18)
Time (months)
b
% of tumors with Gain/loss
S1 (older)
S2
S3(youngest)
c
% of tumors with Gain/loss
C1
C2
C3
C4
C5
C6
d
C4 Subset :Survival based on Gender
Percent survival
Female
p = 0.015
Male
Time (months)
C4 Subset
e
C4 Subset :Survival based on CN for 4p
Percent survival
Copy# Deletions
p = 0.032
Copy# neutral
Time (months)
f
Wound healing
Complement Activation
Metabolic Process
Runx2 regulates chondrocyte maturation
Runx1 regulates transcription in IL signaling
Nonsense Mediated decay

Supp Fig 4

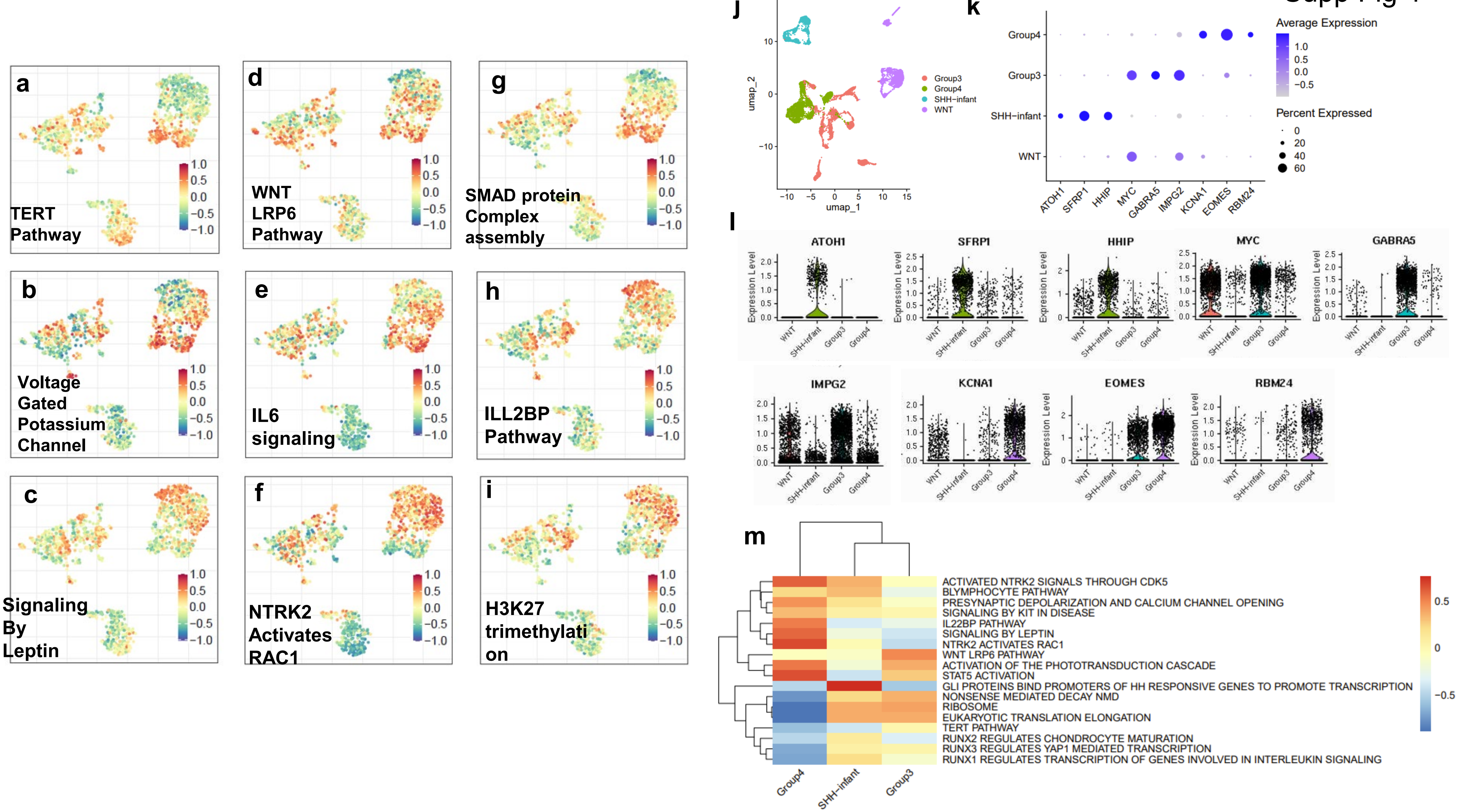
a
TERT Pathway
b
Voltage Gated Potassium Channel
c
Signaling By Leptin
d
WNT LRP6 Pathway
e
IL6 signaling
f
NTRK2 Activates RAC1
g
SMAD protein Complex assembly
h
ILL2BP Pathway
i
H3K27 trimethylati on
j
Group3
Group4
SHH−infant
WNT
umap_1
umap_2
k
Average Expression
Percent Expressed
Group4
Group3
SHH−infant
WNT
ATOH1
SFRP1
HHIP
MYC
GABRA5
IMPG2
KCNA1
EOMES
RBM24
l
ATOH1
SFRP1
HHIP
MYC
GABRA5
IMPG2
KCNA1
EOMES
RBM24
Expression Level
m
ACTIVATED NTRK2 SIGNALS THROUGH CDK5
BLYMPHOCYTE PATHWAY
PRESYNAPTIC DEPOLARIZATION AND CALCIUM CHANNEL OPENING
SIGNALING BY KIT IN DISEASE
IL22BP PATHWAY
SIGNALING BY LEPTIN
NTRK2 ACTIVATES RAC1
WNT LRP6 PATHWAY
ACTIVATION OF THE PHOTOTRANSDUCTION CASCADE
STAT5 ACTIVATION
GLI PROTEINS BIND PROMOTERS OF HH RESPONSIVE GENES TO PROMOTE TRANSCRIPTION
NONSENSE MEDIATED DECAY NMD
RIBOSOME
EUKARYOTIC TRANSLATION ELONGATION
TERT PATHWAY
RUNX2 REGULATES CHONDROCYTE MATURATION
RUNX3 REGULATES YAP1 MEDIATED TRANSCRIPTION
RUNX1 REGULATES TRANSCRIPTION OF GENES INVOLVED IN INTERLEUKIN SIGNALING
Group4
SHH−infant
Group3

Supp Fig 5

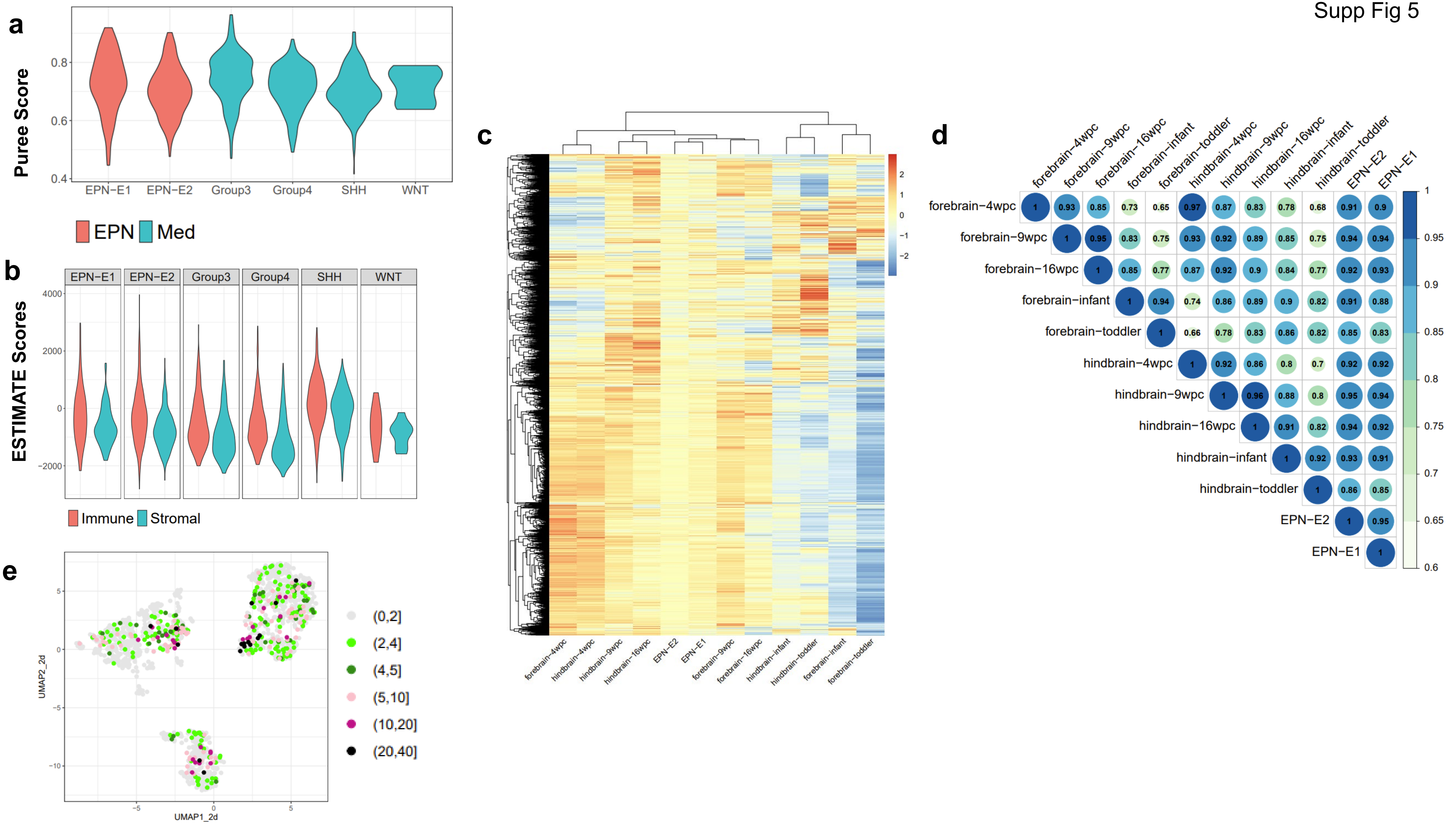
a
Puree Score
0.4
0.6
0.8
EPN−E1
EPN−E2
Group3
Group4
SHH
WNT
EPN
Med
b
ESTIMATE Scores
EPN−E1
EPN−E2
Group3
Group4
SHH
WNT
4000
2000
0
−2000
Immune
Stromal
c
2
1
0
−1
−2
forebrain−4wpc
hindbrain−4wpc
hindbrain−9wpc
hindbrain−16wpc
EPN−E2
EPN−E1
forebrain−9wpc
forebrain−16wpc
hindbrain−infant
hindbrain−toddler
forebrain−infant
forebrain−toddler
d
forebrain−4wpc
forebrain−9wpc
forebrain−16wpc
forebrain−infant
forebrain−toddler
hindbrain−4wpc
hindbrain−9wpc
hindbrain−16wpc
hindbrain−infant
hindbrain−toddler
EPN−E2
EPN−E1
forebrain−4wpc 1 0.93 0.85 0.73 0.65 0.97 0.87 0.83 0.78 0.68 0.91 0.9
forebrain−9wpc 1 0.95 0.83 0.75 0.93 0.92 0.89 0.85 0.75 0.94 0.94
forebrain−16wpc 1 0.85 0.77 0.87 0.92 0.9 0.84 0.77 0.92 0.93
forebrain−infant 1 0.94 0.74 0.86 0.89 0.9 0.82 0.91 0.88
forebrain−toddler 1 0.66 0.78 0.83 0.86 0.82 0.85 0.83
hindbrain−4wpc 1 0.92 0.86 0.8 0.7 0.92 0.92
hindbrain−9wpc 1 0.96 0.88 0.8 0.95 0.94
hindbrain−16wpc 1 0.91 0.82 0.94 0.92
hindbrain−infant 1 0.92 0.93 0.91
hindbrain−toddler 1 0.86 0.85
EPN−E2 1 0.95
EPN−E1 1
1
0.95
0.9
0.85
0.8
0.75
0.7
0.65
0.6
e
UMAP2_2d
UMAP1_2d
5
0
−5
−10
(0,2]
(2,4]
(4,5]
(5,10]
(10,20]
(20,40]

Supp Fig 6

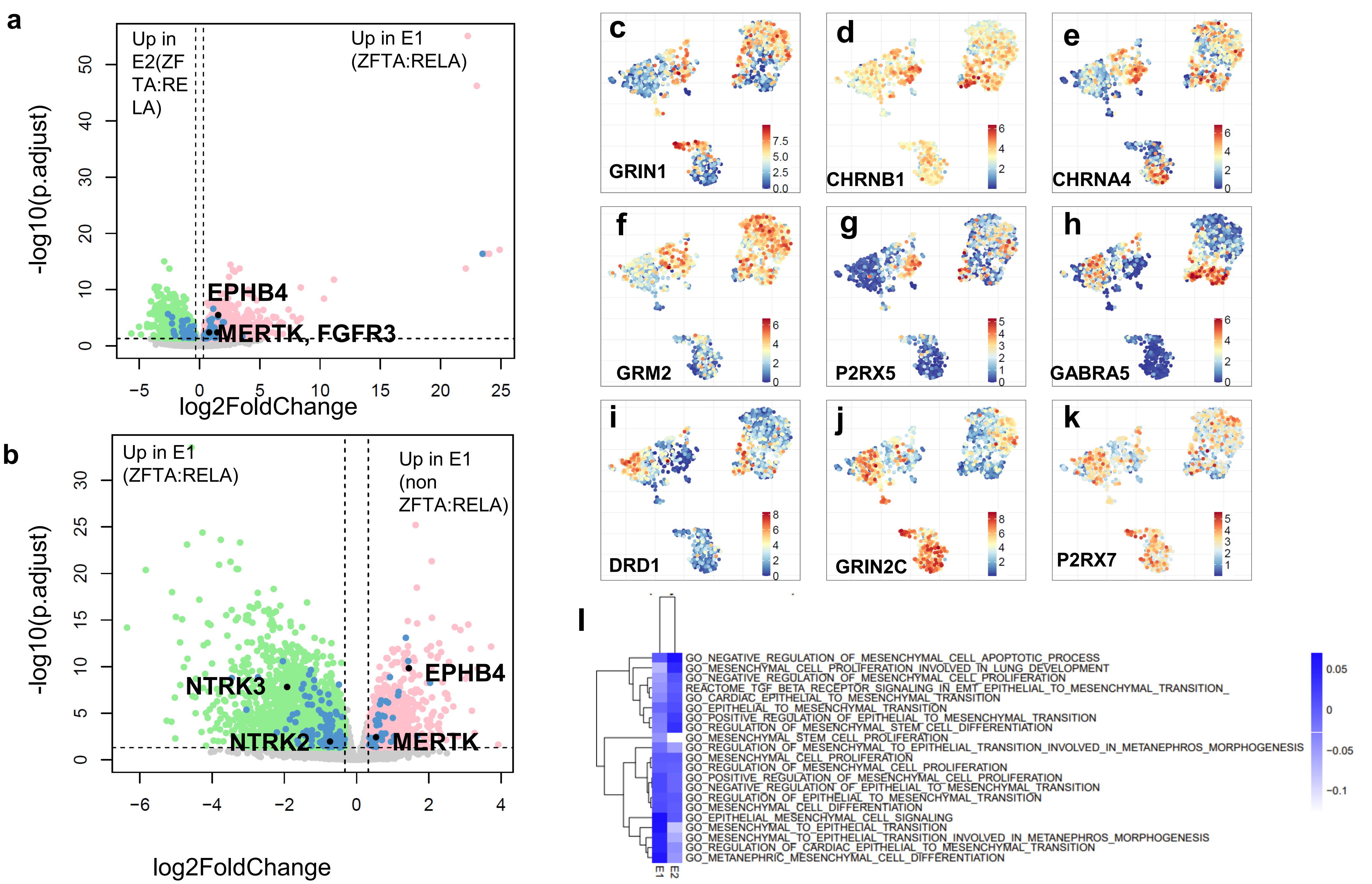
a
Up in E2(ZFTA:RELA)
Up in E1 (ZFTA:RELA)
EPHB4
MERTK, FGFR3
-log10(p.adjust)
log2FoldChange
b
Up in E1 (ZFTA:RELA)
Up in E1 (non ZFTA:RELA)
EPHB4
NTRK3
NTRK2
MERTK
-log10(p.adjust)
log2FoldChange
c
GRIN1
d
CHRNB1
e
CHRNA4
f
GRM2
g
P2RX5
h
GABRA5
i
DRD1
j
GRIN2C
k
P2RX7
l
GO_NEGATIVE_REGULATION_OF_MESENCHYMAL_CELL_APOPTOTIC_PROCESS
GO_MESENCHYMAL_CELL_PROLIFERATION_INVOLVED_IN_LUNG_DEVELOPMENT
GO_NEGATIVE_REGULATION_OF_MESENCHYMAL_CELL_PROLIFERATION
REACTOME_TGF_BETA_RECEPTOR_SIGNALING_IN_EMT_EPITHELIAL_TO_MESENCHYMAL_TRANSITION_
GO_CARDIAC_EPITHELIAL_TO_MESENCHYMAL_TRANSITION
GO_EPITHELIAL_TO_MESENCHYMAL_TRANSITION
GO_POSITIVE_REGULATION_OF_EPITHELIAL_TO_MESENCHYMAL_TRANSITION
GO_REGULATION_OF_MESENCHYMAL_STEM_CELL_DIFFERENTIATION
GO_MESENCHYMAL_STEM_CELL_PROLIFERATION
GO_REGULATION_OF_MESENCHYMAL_TO_EPITHELIAL_TRANSITION_INVOLVED_IN_METANEPHROS_MORPHOGENESIS
GO_MESENCHYMAL_CELL_PROLIFERATION
GO_REGULATION_OF_MESENCHYMAL_CELL_PROLIFERATION
GO_POSITIVE_REGULATION_OF_MESENCHYMAL_CELL_PROLIFERATION
GO_NEGATIVE_REGULATION_OF_EPITHELIAL_TO_MESENCHYMAL_TRANSITION
GO_REGULATION_OF_EPITHELIAL_TO_MESENCHYMAL_TRANSITION
GO_MESENCHYMAL_CELL_DIFFERENTIATION
GO_EPITHELIAL_MESENCHYMAL_CELL_SIGNALING
GO_MESENCHYMAL_TO_EPITHELIAL_TRANSITION
GO_MESENCHYMAL_TO_EPITHELIAL_TRANSITION_INVOLVED_IN_METANEPHROS_MORPHOGENESIS
GO_REGULATION_OF_CARDIAC_EPITHELIAL_TO_MESENCHYMAL_TRANSITION
GO_METANEPHRIC_MESENCHYMAL_CELL_DIFFERENTIATION
E1
E2
0.05
0
-0.05
-0.1

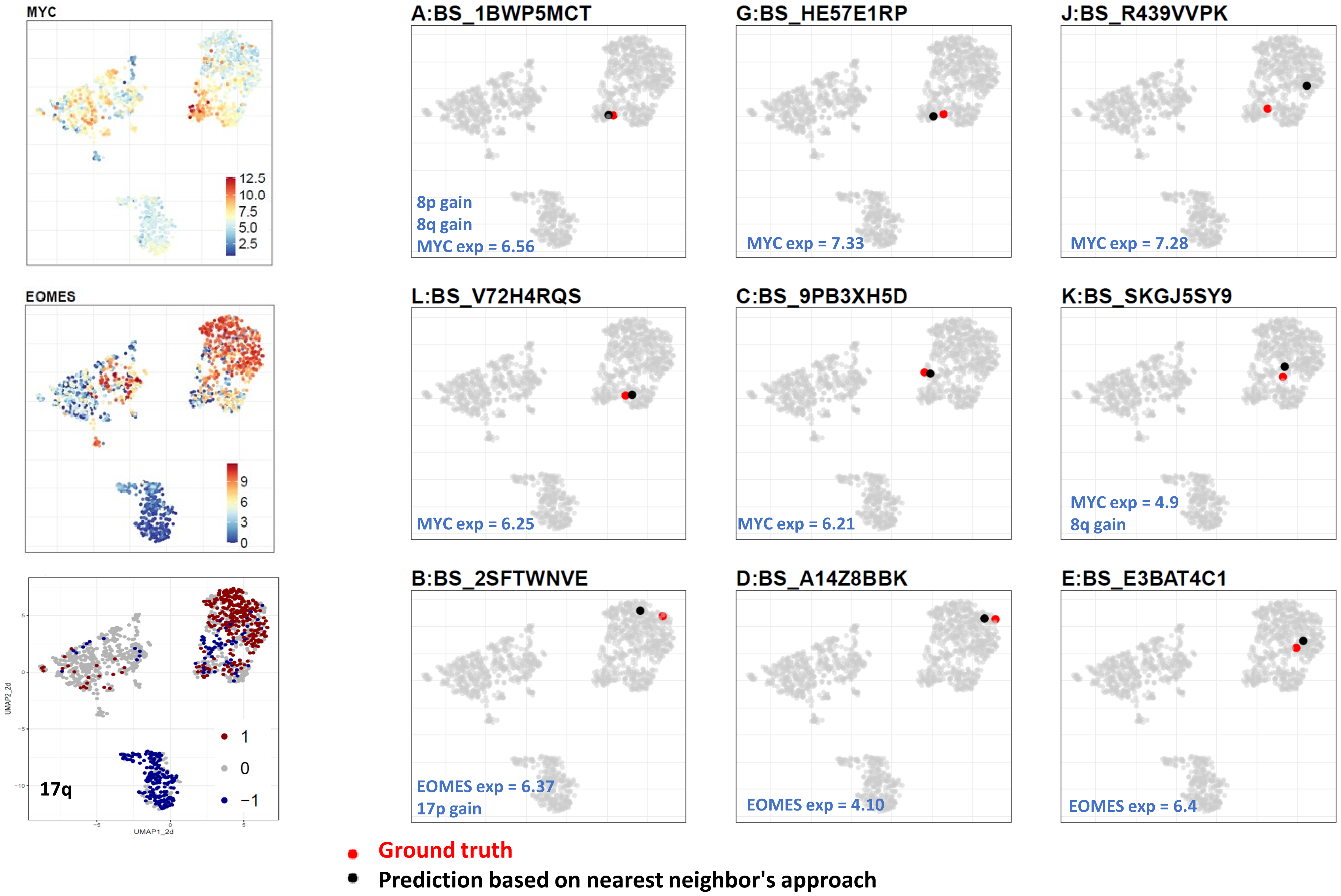
MYC
12.5
10.0
7.5
5.0
2.5
EOMES
9
6
3
0
17q
1
0
−1
UMAP1_2d
UMAP2_2d
A:BS_1BWP5MCT
8p gain
8q gain
MYC exp = 6.56
G:BS_HE57E1RP
MYC exp = 7.33
J:BS_R439VVPK
MYC exp = 7.28
L:BS_V72H4RQS
MYC exp = 6.25
C:BS_9PB3XH5D
MYC exp = 6.21
K:BS_SKGJ5SY9
MYC exp = 4.9
8q gain
B:BS_2SFTWNVE
EOMES exp = 6.37
17p gain
D:BS_A14Z8BBK
EOMES exp = 4.10
E:BS_E3BAT4C1
EOMES exp = 6.4
Ground truth
Prediction based on nearest neighbor's approach

### Suppl Figure legends

**Fig S1. Dimension Reduction and Normalization Techniques Applied to RNA-seq Data** (A-C) Dimension reduction techniques applied to non-batch corrected log2(TPM+1) data: (A) PCA, (B) t-SNE, (C) UMAP. (D-F) Dimension reduction techniques applied to batch-corrected log2(TPM+1) data: (D) PCA, (E) t-SNE, (F) UMAP. (G-I) Different normalization methods applied to batch-corrected data: (G) CPM normalization, (H) RPKM normalization, (I) VST normalization. (J) Stacked barplots showing proportion of samples with amplifications and deletions in chr arm 17p, 17q, 1q, 22q, 9p, 9q (K) Boxplots showing gene expression ( log2(TPM+1)) of genes : ATOH1, MYC, KCNA1, WNT5A , TGFB1 and IGF2.

**Fig S2. Further Validation of the Reference Landscape and Subtypes of Medulloblastoma and Ependymoma** (A-F) Validation based on gene expression patterns across different medulloblastoma and ependymoma subtypes. SFRP1 and HHIP show elevated expression of SHH medulloblastoma, GABRA5 and IMPG2 show elevlates expression in Group3 medulloblastoma whereas EOMES and RBM24 show elevated gene expression in Group4 medulloblastoma patients. (G-K) Validation based on gene fusions identified in the different medulloblastoma subtypes, as previously shown by Luo et al[23].

**Fig S3. Copy Number analysis for various medulloblastoma clusters** (A) Survival analysis for S2 based on the copy number profile of 9q. (B) Manhattan plot showing the percentage of tumors with amplifications and deletions in each of the 3 subclusters of Shh medulloblastoma - S1, S2 and S3 and(C) six clusters of group3 and group4 medulloblastoma. (D=E) Kaplan-Meier plots for C4 based on gender and copy number status of 4p (Male(blue) vs Female(pink), p=0.015, n[Male]=18, n[Female]=6, Copy No deletions(green) vs Copy no neutral(grey), pvalue =0.032, n[Copy no deletions]=10, n[copy no neutral] =14).

**Fig S4. Clustering Methods Validate Group 3 and Group 4 Medulloblastoma Clusters** (A-I) GSVA scores for pathways regulated in Group 3 and Group 4 medulloblastoma, visualized on the UMAP. (J) UMAP showing scRNASeq data from 25 patients colored in subtype (K) Dotplot showing gene expression of genes (x-axis) in each of subtypes for scRNASeq data (L) ViolinPlots showing gene expression of genes (x-axis) in each of subtypes for scRNASeq data (M) Heatmap showing the average pathway score for pathways shown in Fig4, for each subtype of scRNASeq data.

**Fig S5. Clustering Methods Validate EPN-E1 and EPN-E2**

Tumor purity estimates based on (A) PurEE and (B) ESTIMATE. (C) Heatmap displaying the average expression of the 8,000 most variable genes across forebrain and hindbrain samples, as well as EPN-E1 and EPN-E2 samples. (D) Correlation plot illustrating relationships among forebrain and hindbrain samples and EPN-E1 and EPN-E2, based on the 8,000 most variable genes. (E) Gene fusion frequencies across all ependymoma and medulloblastoma samples.

**Fig S6. Differential gene expression analysis for EPN-E1 and EPN-E2** (A) Volcano plot showing DEGs upregulated in E1 containing ZFTA:RELA gene fusions vs E2 containing ZFTA:RELA gene fusions. (b) Volcano plot showing DEGs upregulated within E1 containing ZFTA:RELA vs those that did not contain ZFTA:RELA gene fusion (C-K) Synaptic genes upregulated in EPN -E1 and EPN-e2 respectively (L) Heatmap showing average of mesenchymal scores across EPN-E1 and EPN-E2.

**Fig S7. Validation of New Patient Data Overlay on the Reference Landscape** Validation of a new patient overlayed on the reference landscape, based on gene expression and copy number patterns.

### Suppl Tables

- **Table S1**. Datasets from North America and Europe were combined to generate the medulloblastoma UMAP

- **Table2 S2** Copy number profiles for each subtype of Group3 and Group4 medulloblastoma , and each syubtype of shh medulloblastoma, as obtained from consensus clustering.

- **Table S3** Top recurrent gene fusions in ST-EPNs in EPN-E1

- **Table S4** Top recurrent gene fusions in PF-EPNs in EPN-E2

- **Table S5** Top recurrent gene fusions in medulloblastoma subtypes

- **Table S6** All genes and kinases upregulated in EPN-E1 and EPN=E2

- **Table S7** Pathways upregulated in EPN-E1 and EPN-E2

- **Table S8** Mapping of 12 NOS medulloblastoma samples from CBTN as shown in Figure 7

.
